# Degradation of LMO2 in T cell leukaemia results in collateral breakdown of transcription complex partners and causes LMO2-dependent apoptosis

**DOI:** 10.1101/2024.12.09.627495

**Authors:** Naphannop Sereesongsaeng, Carole J.R. Bataille, Angela J. Russell, Nicolas Bery, Fernando Sialana, Jyoti Choudhary, Ami Miller, Terry H. Rabbitts

**Author notes:** Nicolas Bery: Université de Toulouse, Inserm, CNRS, Université Toulouse III-Paul Sabatier, Centre de Recherches en Cancérologie de Toulouse, 31100 Toulouse, France.; Ami Miller: Evotec,114 Innovation Dr., Milton Park, Abingdon, OX14 4RZ, UK. Corresponding author: Terry Rabbitts.

## Abstract

LMO2 is an intrinsically disordered transcription factor activated in T cell leukaemia that is difficult to target. It forms part of a multiprotein complex that has bipartite DNA binding through heterodimeric bHLH and GATA proteins. To determine if degradation of LMO2 in the context of T-ALL has therapeutic potential, a chimaeric intracellular antibody has been developed fusing an anti-LMO2 single domain variable region with one of three E3 ligases to create biodegraders. The intracellular binary interaction of these biodegraders with LMO2 leads to its proteosomal degradation but, in addition, concomitant loss of bHLH proteins that associate with LMO2 in the DNA-binding complex. Chemical compound surrogates of the intracellular antibody paratope (called Abd compounds) have been modified to create proteolysis targeting chimaeras (PROTACs) for orthogonal assays of effects of LMO2 degradation. These form a ternary complex with LMO2 and E3 ligase in leukaemia cells that induces degradation of LMO2, and is also accompanied by loss of associated bHLH proteins. This is accompanied by T-ALL growth inhibition, alterations in proteins involved in cell cycling and instigation of apoptosis. These effects do not occur in the absence of LMO2. Our work demonstrates that degradation of LMO2 affects T-ALL and the lead compounds can eventually be developed into drugs for patient treatment. Our work describes methods for drug discovery starting with antibody fragments.

## INTRODUCTION

Chromosomal translocations are among the consistent somatic mutations that lead to cancer and to the maintenance of the malignancy by abnormal gene activation and/or creating fusion genes https://mitelmandatabase.isb-cgc.org/ (1, 2). These changes are found in all tumour types and regulation of the tumour state is due to the translocation gene expressed proteins. In tumours of haematopoietic origin and some solid tumours, for instance, sarcomas the translocation proteins are transcription factors that can have master regulator functions in tumourigenesis by disrupting differentiation programmes (3, 4). While translocation proteins offer tumour-specific targets and therefore attractive as therapeutic targets, where they are transcription factors, they are considered undruggable, or hard-to-drug. This is because they are intrinsically disordered proteins or have extensive disordered regions that makes them difficult to handle in recombinant form and for which limited structural information are known.

In T cell acute leukaemias (T-ALL), there are number of chromosomal translocations (5, 6), including those activating the *LMO2* gene by t(11;14)(p13;q11) (7, 8). The LMO2 protein is a master regulator of haematopoiesis since mouse gene targeting showed that both primitive (9) and definitive haematopoiesis fail with loss of the *Lmo2* gene (10). Further, this regulator function of LMO2 is shown by its role in remodelling of established vascular endothelial but not in *de novo* endothelial formation (11, 12). This latter function has implication for targeting tumour angiogenesis. LMO2 is a transcription factor whose role in creating a multi-protein complex was first demonstrated in erythroid cells and the LMO2 complex comprises LDB1, TAL1, E47 and GATA (13). Although LMO2 is a zinc containing protein comprising to LIM domains, each with two LIM fingers, the protein does not appear to interact directly with DNA but rather bridges a multi-protein DNA-binding complex (14).

LMO2 is an intrinsically disordered protein (IDP) that defied production in recombinant form and it has been hard to study the cellular properties of the protein and the protein complex. We have described anti-LMO2 human single variable **i**ntracellular **<u>D</u>**omain **<u>A</u>**nti**<u>b</u>**ody (designed **iDAb**) that blocks the formation of the LMO2 protein complex when expressed in cells (15) and discriminates the LMO family paralogues (16). In addition, we used a cell-based intracellular antibody competition assay to develop compounds to interfere with the LMO2-iDAb interaction and these small molecules that are surrogates of the iDAb paratope (17) (**<u>A</u>**nti**<u>b</u>**ody **<u>d</u>**erived (**Abd**) compounds). Further, chimaeric intracellular antibodies could be engineered to carry warheads such as pro-caspase that cause proximity-induced, antigen-dependent apoptosis (18) and suggested that other fusion proteins could be designed to affect other cellular pathways such as proteolysis (19). This was demonstrated using our anti-RAS iDAb for targeted protein degradation of KRAS (20). As a means to study the LMO2 transcription complex in the context of T cell acute leukaemia, we have produced two types of protein degradation mediators. In the first, an anti-LMO2 iDAb was directly fused with one of three different E3 ligases (herein called biodegraders) and in the second, an LMO2 iDAb paratope derived Abd compound was altered to be two different Proteolysis Targeting Chimaeras (PROTACs) (21–24). Treatment of T-ALL cells expressing LMO2, from chromosomal translocations or other promoter alterations, with these degraders showed rapid targeted protein degradation of the LMO2 transcription factor but also levels of TAL1 and E47 (members of the transcription complex (13)) are reduced. These alterations in the LMO2 multi-protein complex cause inhibition of cell growth accompanied by initiation of programmed cell death. Therefore, targeting protein stability of the LMO2 complex shows the dependency of LMO2-positive T-ALL and that other proteins in the transcription factor complex can be disrupted by collateral degradation. LMO2 degraders are lead compounds for drug development for LMO2-positive T-ALL and the approach is applicable to transcription factor complexes, including those involving IDPs.

## RESULTS

### A biodegrader causes loss of LMO2 and collateral loss TAL1/E47 bHLH

Our mouse models of LMO2 tumourigenesis we observed that transgenic enforced thymus expression of Lmo2 (using CD2 promoter) but not Tal1 causes later onset, clonal T cell neoplasia, which arose earlier in double transgenic mice (25). Expressing anti-LMO2 iDAb (a human VH segment) inhibited tumour growth in a transplantation model (15). To assess the ability to degrade LMO2 by the ubiquitin proteasome system, we begun to dissect the elements of interaction in the LMO2 complex by engineering chimaeric intracellular antibodies where an E3 ligase was fused either at the N- or C-terminus of the iDAb to mediate binary interactions between iDAb-E3 ligase and LMO2 for ubiquitination of LMO2 and proteasomal degradation. Targeted protein degradation of LMO2 was observed when any of the six iDAb-E3 ligase proteins were expressed in HEK293T cells along with an LMO2 expressing plasmid (Supplementary Fig. 1, panels A, B, D, E, G and H). No detectable loss of control protein RAS was observed (or of the loading controls, cyclophilin-b and β-actin). Conversely, using anti-RAS iDAb-CRBN or anti-KRAS DARPin-UBOX or anti-KRAS DARPin-VHL (Supplementary Fig. 1, panels C, F and I, respectively) caused turnover of RAS but not LMO2.

We further investigated the LMO2 degradation in natural LMO2 expression settings (i.e. T-ALL lines) using a lentivirus that conditionally expresses anti-LMO2 iDAb VH576 fused to CRBN E3 ligase (designated TLCV2-iDAb-CRBN) via a doxycycline-inducible promoter (TLCV2 (26)). CRBN was chosen in the inducible assay since thalidomide is commonly used in PROTAC chemistry and this gave the opportunity to further evaluate LMO2 degradation by cereblon. This lentivirus was used to infect either Jurkat (lacking LMO2 expression), KOPT-K1 (27) or P12-Ichikawa cells (28) (expressing LMO2 after chromosomal translocations). The level of LMO2 expression after induction of the biodegrader was analysed by Western blotting. LMO2 levels showed a marked decrease within 9 hours of iDAb-CRBN induction and continued for the 48 hours of analysis in LMO2-expressing cells (Fig. 1A).

**Figure 1.**
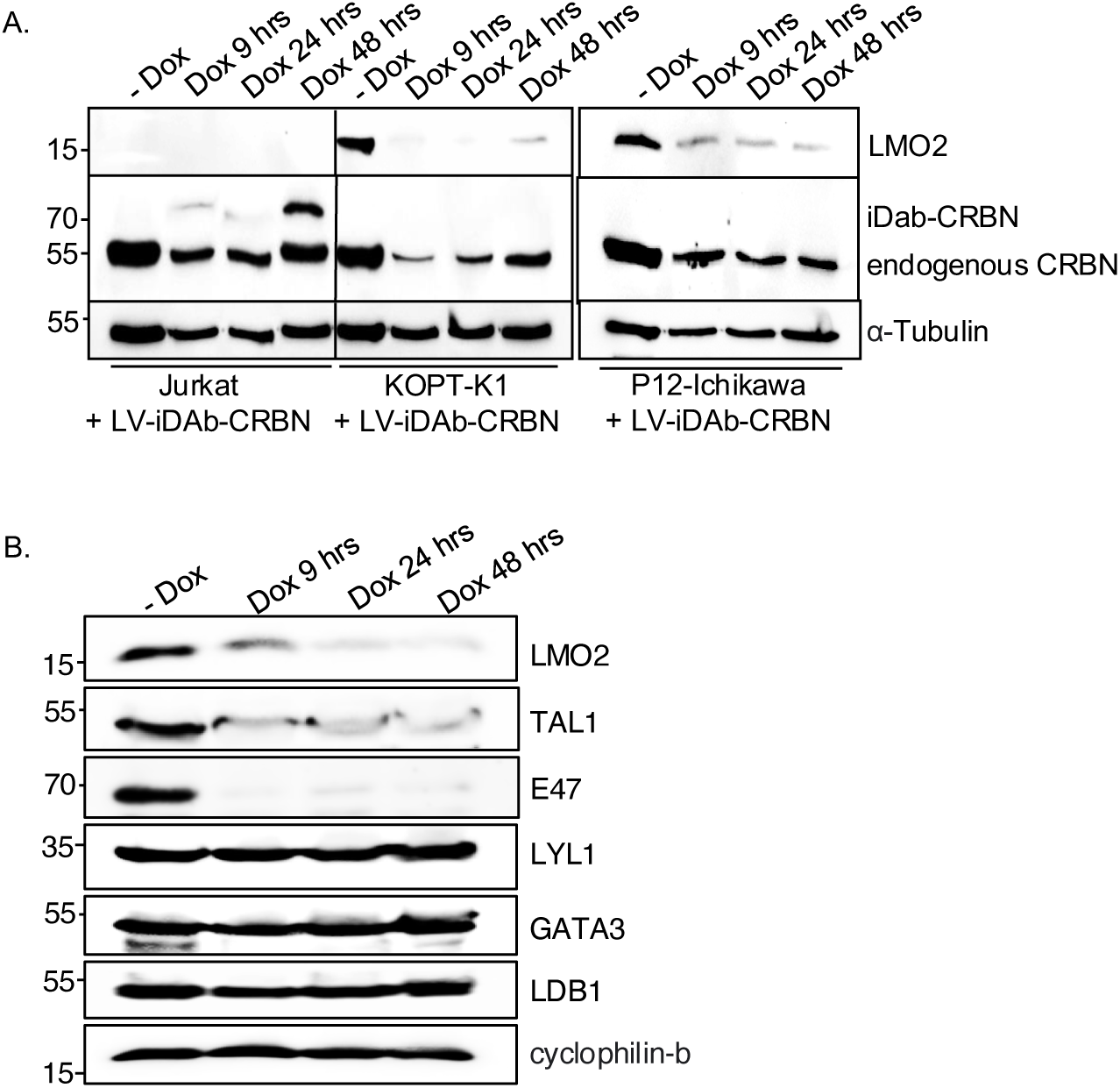
Lentiviral expressed biodegrader affects the LMO2 multi-protein complex in T cell lines. The T cell lines Jurkat (LMO2-), KOPT-K1 (LMO2+), and P12-Ichikawa (LMO2+) cells were infected with lentivirus packaging plasmids and transfer vector (TLCV2-VH576-L10-CRBN) for 16 hours followed by 2 μg/ml doxycycline induction for 9, 24, and 48 hours. Western blotting analysis was used to detect LMO2, endogenous CRBN, and VH576-L10-CRBN levels after lentivirus infection. α-tubulin was used as an internal loading control for Western blotting analysis (panel **A**). The level of LMO2 and proteins associated with the LMO2 transcription complex (TAL1, E47, Lyl-1, GATA3, and LDB1) was determined after KOPT-K1 cells were infected with TLCV2-VH576-L10-CRBN followed by 2 μg/ml doxycycline induction for 9, 24, and 48 hours (panel **B**). Cyclophilin-b were used as an internal loading control.

The LMO2 protein is associated with a bipartite DNA-binding multi-protein complex, involving basic-helix-loop-helix (bHLH) proteins TAL1/SCL and E47 and GATA (13). We determined if other members of the complex could be degraded coincidentally with LMO2 using the anti-LMO2 biodegrader. KOPT-K1 cells, which express LMO2 from a t(11;14)(p13:q11) chromosomal translocation (27), was infected with the lentiviral TLCV2-iDab-CRBN and the biodegrader expression was induced for 9, 24 and 48 hours with doxycycline. Protein extracts were analysed by Western blotting using antibodies recognizing various components of the LMO2 complex. Targeted protein degradation of LMO2 is observed throughout the course of the biodegrader induction and we also observed a rapid degradation the two associated bHLH proteins TAL1/SCL and E47. No evidence was found for associated turnover of another bHLH protein LYL1 or of the LMO2-associating protein LDB1 or GATA3 (Fig. 1B). This unexpected finding of the bHLH collateral degradation is unlikely to be due to off target binding of the anti-LMO2 iDAb with TAL1 or E47.

### Control of LMO2 protein levels using chemical degraders

These results with the biodegraders demonstrate the degradability of endogenous LMO2 by E3 ligase-based molecules. The difficulty of the delivery of these macromolecules supported the subsequent development of small molecule-based LMO2 degraders. To circumvent the problem of antibody delivery, we have previously developed a technology approach which uses antibody paratopes to select small molecule surrogates (**<u>A</u>**nti**<u>b</u>**ody **<u>d</u>**erived technology, **Abd**) (29). Employing this technology, an Abd chemical series that binds to LMO2 was identified that interferes with the protein complex in cells (17). As a means to compare their cell-specific effects with the biodegraders, the Abd chemical compounds have been reformatted as PROTAC Abd degraders (30). We used Abd-L21 and Abd-L27 as starting points for the PROTAC design (Supplementary Fig. 2A) based on previously described SAR (17) where a broad tolerance of a range of para-substituents on the right-hand side arene ring was found (Fig. 2A). Similarly on the left-hand side, meta-substituted arenes were preferred with a range of substituents tolerated. Piperazine was preferred to the imidazolidin-2-one, seen in Abd-L9 (Supplementary Fig. 2A), for activity, stability, and ease of synthesis, and was therefore selected for our PROTACs. Thiazole and oxazole were found to be interchangeable without any noticeable difference in activity, although the thiazole series were found to be easier to synthesize with higher chemical stability and better yields. It was decided to use a medium length linker as a starting point, namely two PEG units for the VHL-binding ligand (31) and a six carbon alkyl unit for the CRBN-binding ligand (32). Accordingly, two Abd E3 ligase ligand hetero-bifunctional molecules, designated Abd-VHL and Abd-CRBN, were synthesized with either the VHL or thalidomide E3 ligase ligand respectively (Fig. 2B, C).

**Figure 2.**
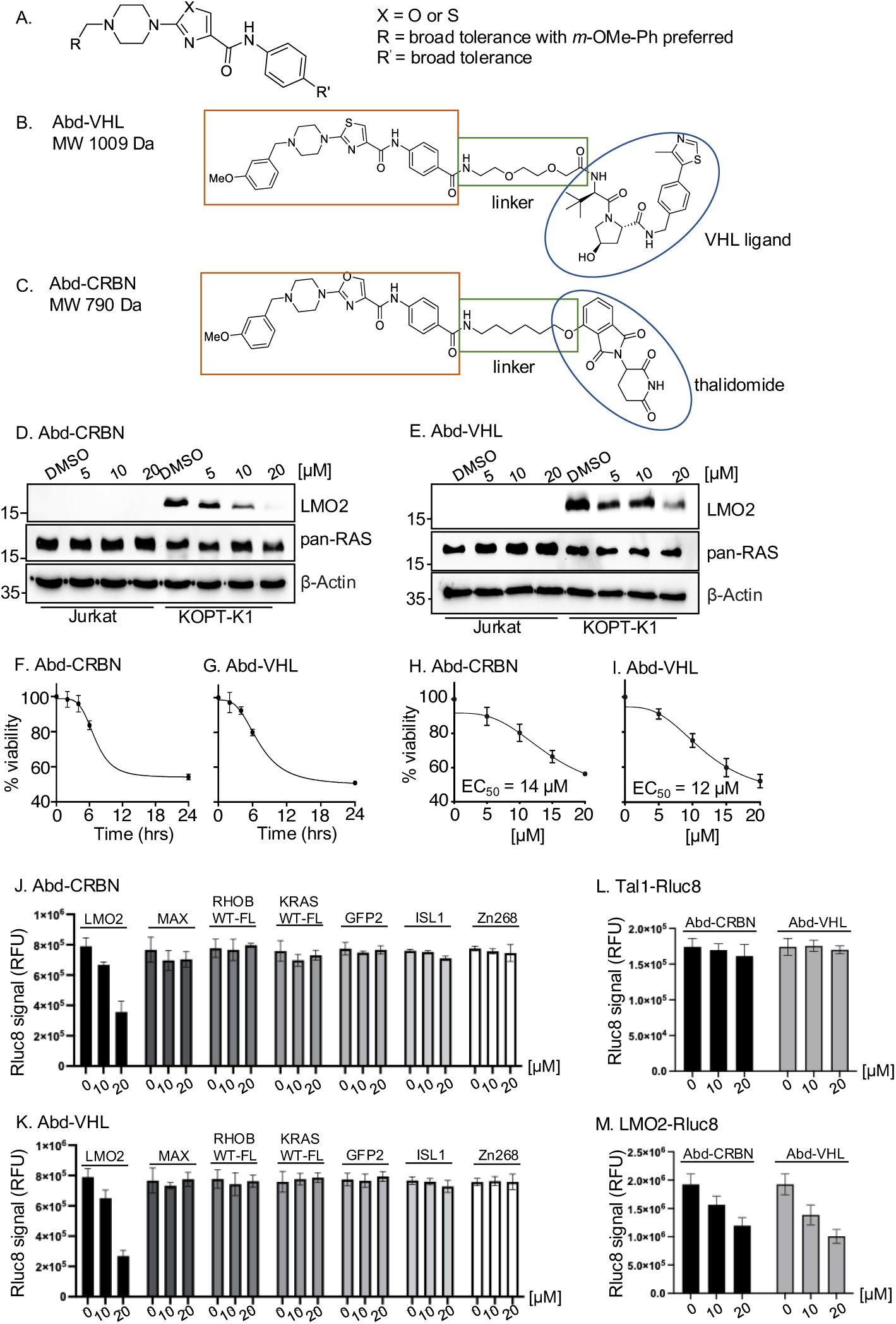
LMO2 Abd degraders and effects on cellular protein. Structure-activity observations of LMO2-binding compounds derived using iDAb VH576 in competitive, cell-based BRET (designated Abd compounds) contributed to a general structure of the LMO2-binding compounds (panel **A**). Based on broad tolerance to chemical modification at position R’, two PROTACs were synthesised, one bearing the VHL ligand (Abd-VHL, panel **B**) and the other bearing thalidomide (the CRBN ligand) (Abd-CRBN, panel **C**). Each PROTAC thus comprises the LMO2-binding ligand, a linker and an E3 ligase ligand. KOPT-K1 and Jurkat cells were treated with different concentration of Abd-degraders at 0, 5, 10, and 20 μM for 24 hours, protein extracts prepared and fractioned by SDS-PAGE followed by Western blotting with the antibodies indicated by each panels **D and E.** Abd-CRBN was used on cells in panels **D** and Abd-VHL in panels **E**. Cell treated with 1% DMSO was used as a control and β-actin was used as an internal loading control for the Western blotting analysis. The 24 hours Western blot data (panels D, E) for KOPT-K1 treated cells were quantitated using densitometry (Supplementary Fig. S3 G and J) to calculate DC_50_ 9 μM and 15 μM for Abd-VHL and Abd-CRBN respectively. The viability of KOPT-K1 cells were treated with 15 μM Abd-CRBN (panel F) or 15 μM Abd-VHL (panel **G**) for 0, 2, 4, 6, and 24 hours viability determined using CellTiter-Glo assay. Data are presented as a relative to luminescence at 0 hours, normalized to 100%. Determination of EC_50_ values of Abd-CRBN and Abd-VHL after 24 hours treatment in KOPT-K1 was determined using CellTiter-Glo calculated by GraphPad Prism 9.0 software (panel **H and I**, respectively). All the values were presented as the average values relative to cell viability values in control (DMSO treated cells) normalized to 100%. Data represent mean ± SEM (n=3). Luciferase cell-based report assays were used to determine the degradation of the proteins after transfected HEK293T cells with different Renilla luciferase (Rluc8) reporter plasmids (pEF-LMO2-Rluc8, pEF-MAX-Rluc8, pEF-RHOBWT-FL, pEF-KRSWT-Fl-RLuc8, pEF-GFP2-Rluc8, pEF-ISL1-Rluc8, and pEF-Zn268) followed by Abd-CRBN (panel **J**) and Abd-VHL (panel **K**) treatment at 0, 10 and 20 μM for 24 hours. The potential degradation of pEF-Tal-Rluc8 (panel **L**) and pEF-LMO2-Rluc8 (panel **M**) was determined by luciferase cell-based report assays after the treatment with Abd-CRBN or Abd-VHL art 0, 10 and 20 μM for 24 hours. Data represent mean + SEM (n=3). The DC_50_ values for loss of LMO2-Rluc8 signal in panel M were calculated as 10 μM and 9 μM for Abd-VHL and Abd-CRBN respectively.

The selectivity of the chemical degraders was tested by comparing T cell viabilities when treated with Abd-VHL and Abd-CRBN or the starting Abd compounds. The result showed that reformatting the LMO2 Abd compounds as PROTACs improved selectivity of LMO2-expressing cells compared to non-expressor ones (Supplementary Fig. 2B). This difference was clearer when titrating the Abd-VHL and Abd-CRBN compounds over a concentration range during a 24-hour assay period. Jurkat (LMO2 negative) were largely unaffected, while KOPT-K1 (LMO2 positive) showed loss of viability with increased PROTAC concentration (Supplementary Fig. 2C, D), which is an effect sustained over 48 hours (Supplementary Fig. 2E, F).

The effect of the Abd-CRBN or Abd-VHL compounds on LMO2 protein levels was then assessed in dose response in KOPT-K1 and Jurkat cells. Cells were treated with a single dose of the compound, ranging from 0-20 µM for 2, 6 or 24 hours followed by analysis of protein levels by Western blotting analysis (Fig. 2 D, E and Supplementary Fig. 3). At 24 hours treatment, LMO2 levels are reduced by either Abd-CRBN (Fig. 2D) or Abd-VHL (Fig. 2E) whereas control protein RAS levels are unaffected. There is a progressive loss of LMO2 protein from 6 hours onwards up to the end point of 24 hours and, after quantification by densitometry (Supplementary Fig. 3E-J), the DC_50_ values at 24 hours (calculated using the Prism9 GraphPad Software) were 9 μM and 15 μM for Abd-CRBN and Abd-VHL respectively. In addition, the loss of signal of the LMO2-Rluc reporter protein from PROTAC treated cells shown in Figure 2M has been used to calculate half-points. Although not strictly DC_50_ as it measures a reporter protein level, these yielded values are 10 μM for Abd-CRBN and 9 μM Abd-VHL.

These findings were paralleled by loss of cell viability over the 24 hours period when KOPT-K1 cells were treated with 15 μM of either compound (Fig. 2F-I) showing the half-maximal effective concentration (EC_50_) in KOPT-K1 was 14 μM and 12 μM for Abd-CRBN and Abd-VHL, respectively. The EC_50_ values were in a comparable range MOLT-3 cells (another LMO2 expressing cell line) (Supplementary Fig. 4, C, F) compared to Jurkat cells that are resistance to either compound (Supplementary Fig. 4A, D). The longevity of degradation was followed using Western blotting and CellTitre-Glo assays (Supplementary Fig. 5A, B and C, D respectively). LMO2 was lost in KOPT-K1 during the 72 hours of cell culture after a single dose of compounds while little or no effect was observed in the LMO2 non-expressing Jurkat cells range.

The possible effect of the compounds on non-LMO protein controls was assessed using a luciferase cell-based report assay (Fig. 2J, K). Fusions with Renilla luciferase of various baits were expressed in HEK293T cells, including the LIM protein ISL1 and the zinc finger proteins Zn268, and luciferase levels determined after treatment with the PROTAC compounds (0-20 μM) for 24 hours. Only loss of luciferase in the positive control LMO2-Rluc8 fusion was observed. In addition, confirmation that the specific involvement of the Abd-CRBN or Abd-VHL to the proteasome machinery in the LMO2 turnover was confirmed by using either proteosome inhibitors or competing the Abd degraders with the respective free E3 ligase ligands (Supplementary Fig. 6). Furthermore, TAL1 protein was fused to Renilla luciferase to investigate possible cross-reaction of the LMO2 PROTACs compounds with TAL1. After treatment with either of the PROTACS compounds (0-20 μM) for 24 hours, the luciferase level does not reduce in TAL1-Rluc8 (Fig. 2L), whereas a reduction of luciferase signal in the positive control LMO2-Rluc8 fusion was observed (Fig. 2M). This result suggests that the collateral degradation of TAL1/SCL, and of E47, are due to biological effects beyond simple protein-compound interaction, as also found with the application of the biodegrader.

### Loss of LMO2 protein also affects partner proteins in the transcription complex

Lentiviral infection of KOPT-K1 cells expressing an LMO2 biodegrader showed collateral loss of TAL1 and E47 proteins with LMO2 (Fig. 1). The bystander degradation was not evident for other members of the transcription complex. To gain further insights into this finding, the effect of the Abd-CRBN and Abd-VHL compounds was assayed with two LMO2 expressing cell lines (KOPT-K1 and CCRF-CEM) compared with two non-expressors (Jurkat and DND-41) (Fig. 3). A similar pattern of LMO2 degradation was observed in the two expressing lines as we had first found with expressed biodegrader from the lentivirus. Allied to this, the loss of both TAL1 and E47 proteins is dose dependent relationship (Fig. 3A), while there was no detectable loss of LYL1 (another T-ALL implicated bHLH protein), GATA3 or LDB1. Importantly, there was no apparent loss of these proteins in the two LMO2-non-expressing T cell lines. These results suggested that the proteolysis of TAL1 and E47 depended on degradation of LMO2 (Fig. 3B). The loss of LMO2, TAL1 and E47 occurs rapidly as protein loss is observed in little as 2 to 4 hours after treatment (Fig. 3C). The involvement of the proteosome was confirmed by inhibition of Abd-CRBN and Abd-VHL by epoxomicin in both KOPT-K1 and CCRF-CEM (Fig. 3D) and requirement for E3 ligase by inhibition with the respective free ligand.

**Figure 3.**
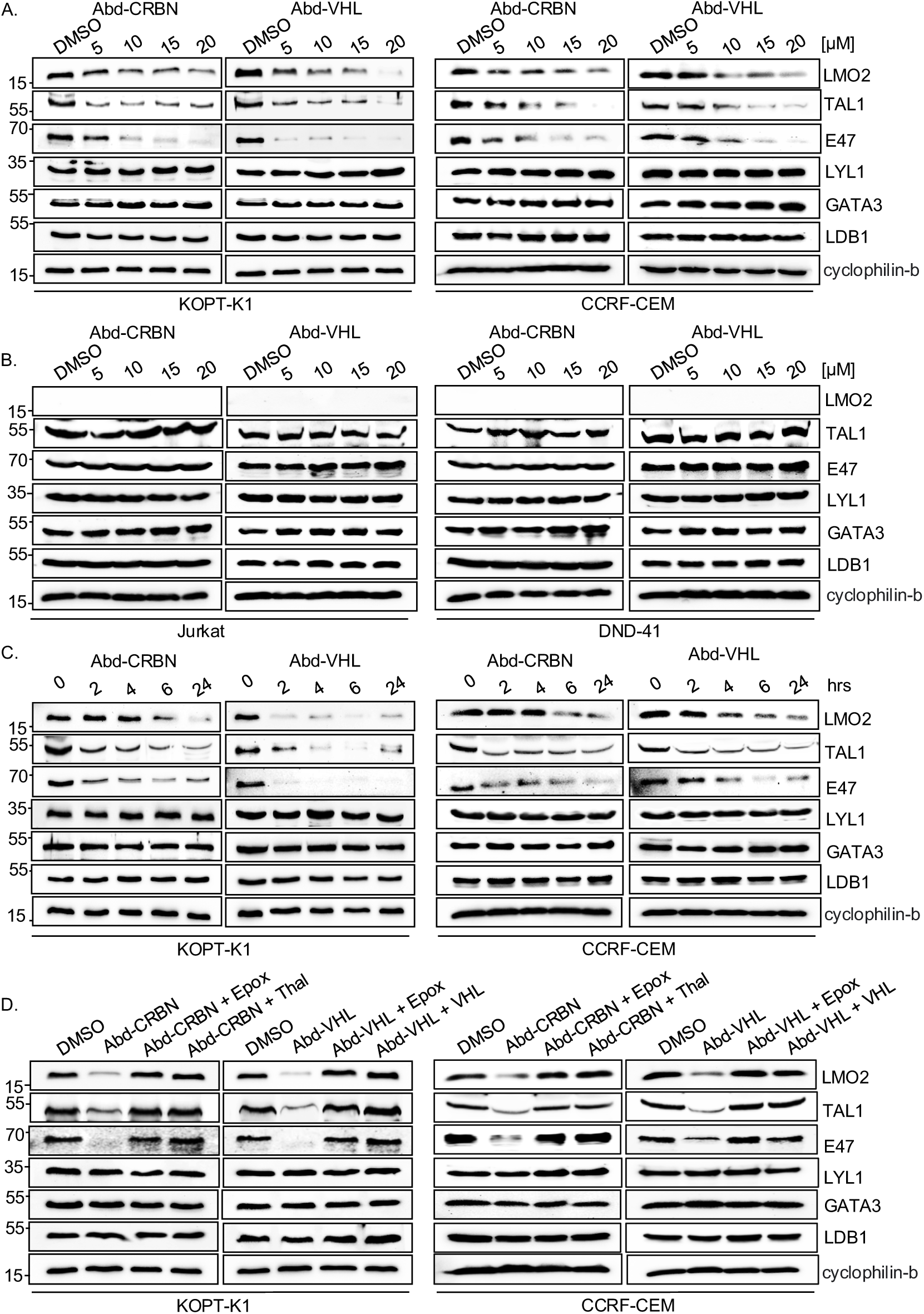
Proteins associated within the LMO2 transcription complex are co-degraded with the LMO2 PROTACs. LMO2 expressing (KOPT-K1 and CCRF-CEM) and non-expressing T cells (Jurkat and DND-41) were treated with Abd-degraders and protein extracts were prepared for Western blot, detecting LMO2, TAL1, E47, Lyl-1, GATA3, LDB1 proteins (cyclophilin-b was used as an internal loading control) after cells were treated with different concentration of Abd-degraders at 0, 5, 10, 15, and 20 μM for 24 hrs. Panel **A**: Western blotting analysis with KOPT-K1 and CCRF-CEM extracts and panel **B** with Jurkat and DND-41 extracts. Panel **C**: KOPT-K1 and CCRF-CEM were treated with Abd-degraders at 20 μM for 2, 4, 6, and 24 hours. Western blotting analysis data show LMO2, TAL1 and E47 proteins expression in KOPT-K1 and CCRF-CEM that was affected by treatment. Panel **D**: The involvement of proteasome machinery in protein complex degradation was investigated in KOPT-K1 and CCRF-CEM with either proteosome inhibitors or by competing the potential of the Abd degraders with free E3 ligase ligand. Western blot data show LMO2, TAL1, and E47 expression in KOPT-K1 and CCRF-CEM after the treatment with or without inhibitors followed by Abd-compounds. Inhibitors used were the proteasome inhibitor epoxomicin (0.8 μM), or CRBN inhibitors (10 μM thalidomide) or free VHL ligand. Cyclophilin-b was used as an internal loading control for Western blotting analysis.

### The degradation of LMO2 interferes with cell growth effecting proteins involved in proliferation

The potential effect of LMO2 protein loss was studied using the panel of cell lines expressing LMO2 (KOPT-K1, PF-382, P12-Ichikawa, CCRF-CEM, MOLT-16 and LOUCY) by simultaneously monitoring protein turnover and cell growth. Cells were treated with a single dose of either LMO2 PROTAC and analysed by Western blot (Fig. 4A-F). The Western blot results showed that LMO2 degradation was observed to have occurred and was sustained for 48 hours. In this time frame we did not find any evidence of loss of RAS proteins or β-actin controls.

**Figure 4.**
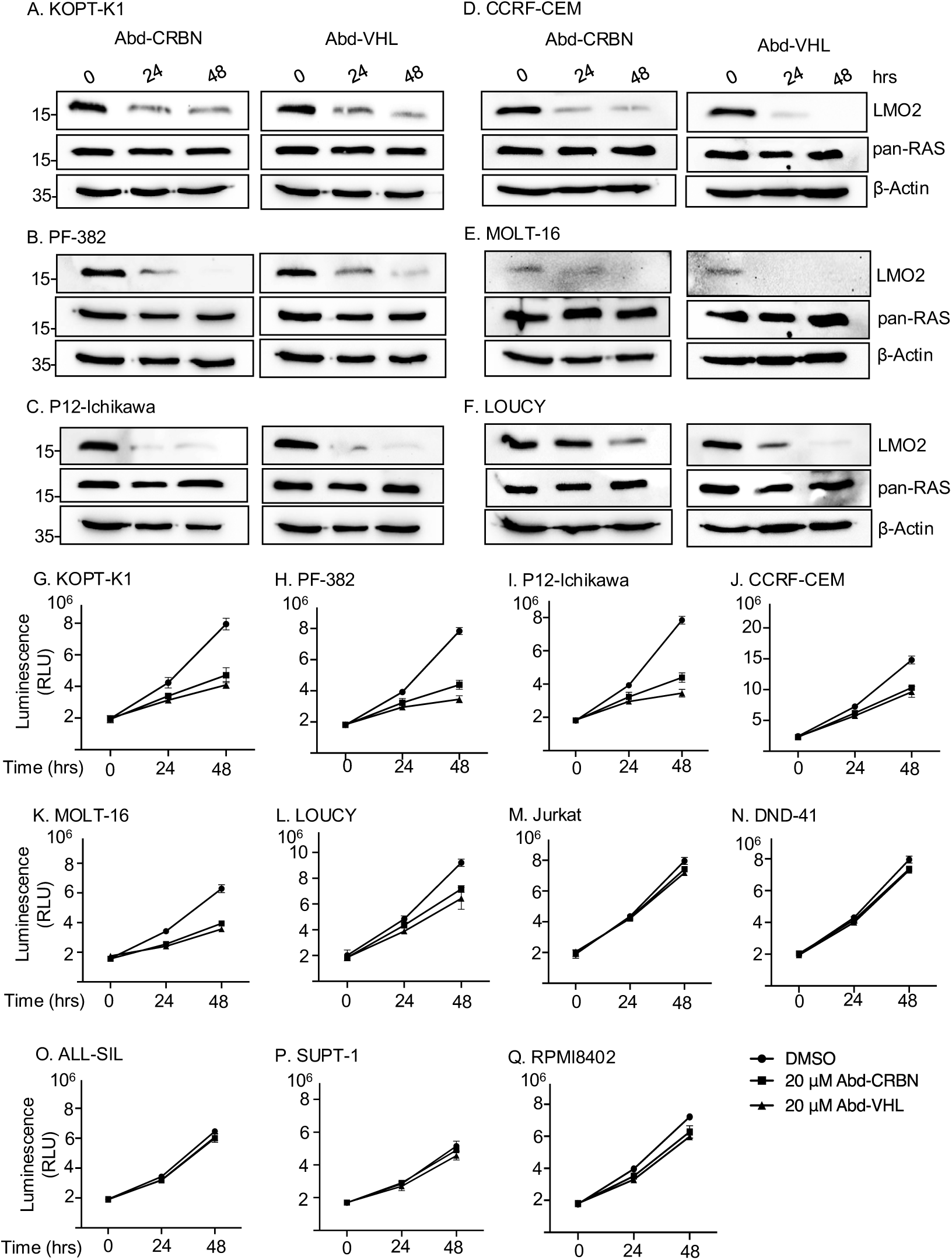
Loss of LMO2 protein inhibits T-ALL cell growth. LMO2 expressing T cells were treated with 20 μM Abd-CRBN or Abd-VHL for 24 and 48 hours and protein extracts were prepared for Western blot to detect LMO2 protein, RAS protein as negative control and β-actin was used as an internal loading control. Western blotting analysis data show LMO2 expression in KOPT-K1 (panel **A**), PF-382 (panel **B**), P12-Ichikawa (panel **C**), CCRF-CEM (panel **D**), MOLT-16 (panel **E**), and LOUCY (panel **F**) that was affected by treatment with the LMO2 PROTAC compounds. Comparative cell numbers were determined from increase in viability determined using the CellTiter-Glo assay. Data represent mean ± SEM (n=3). LMO2 expressing cell lines tested were KOPT-K1 (**G,** t(11;14)(p13;q11)), PF-382 (**H**), P12-Ichikawa (**I**, t(11;14)(p13;q11)), CCRF-CEM (**J**), MOLT-16 (**K**), and LOUCY (**L**). LMO2 non-expressing T cells were also treated with 20 μM Abd-CRBN or Abd-VHL for 24 and 48 hours after which relative luminescence was determined using the CellTiter-Glo assay. Cells tested were Jurkat (panel **M**), DND-41 (panel **N**), ALL-SIL (panel **O**), SUPT-1 (panel **P**), and RPMI8402 (panel **Q**). Data represent mean ± SEM (n=3).

Comparing the effects on growth of cells treated with Abd-CRBN or Abd-VHL, showed an inhibitory effect on T cell growth if LMO2 is expressed (a significant reduction (p<0.001), Fig. 4G-L) whereas T cells without LMO2 (Jurkat, DND-41, ALL-SIL, SUPT-1 and RPMI8402) are largely unaffected (Fig. 4M-Q). The absence of LMO2 protein was confirmed by Western blotting and RT-PCR analysis (Supplementary Fig. 7 and Supplementary Fig. 8, respectively) and those expressing LMO1 was confirmed by RT-PCR (RPMI8402, has an LMO1 chromosomal translocation t(11;14)(p15;q11) from which *LMO1* was cloned (33, 34). The small effects observed at 48 hours may be attributable to off-target toxicity.

Given the short time of exposure to the Abd PROTAC compounds (24 hours) to impair LMO2 positive T-ALL cell growth, we performed proteomic analysis of cells treated with Abd-VHL to ascertain the resultant changes to the cell composition. Proteomic data were obtained in an analysis of Abd-VHL treated in KOPT-K1 (LMO2 expressor), Jurkat (non expressor) and RPMI8402 (non expressor) cells. After the treatment of 15 μM Abd-VHL for 24 hours, the effect on the level of LMO2 was determined using LC-MS with a targeted MS2 method (Fig. 5A) and Western blotting (Fig. 5B), which confirmed the significant reduction of LMO2 after the treatment in KOPT-K1. We collected total proteome dataset and conducted Gene Ontology (GO) and KEGG pathway analyses using rank-based Gene Set Enrichment Analysis (GSEA) across the three cell lines and visualized the resulting enrichment scores in a heatmap (Fig. 5C). Several pathways are affected after the treatment with Abd-VHL, even in the two cell lines that do not express LMO2. Most important, proteins related to cell division appear in the top 10 significantly affected pathways in KOPT-K1 while this effect does not appear in Jurkat and RPMI8402, consistent with cell growth data in KOPT-K1. Further, the average fold change (log_2_FC) in KOPT-K1 (Fig. 5D) and Jurkat (Fig. 5E) was represented as volcano plots, indicating that the total significantly changed proteins in KOPT-K1 is higher than Jurkat cells. To specify the effect of the Abd-VHL treatment to proteins changed in KOPT-K1, the proteins significantly increased or decreased are presented in Fig. 5D and tabulated in Fig. 5F compared to Jurkat cells. The result showed that those significantly down regulated proteins in KOPT-K1 relate to cell division pathway and some aspects of ubiquitination while most proteins in Jurkat cells are unchanged. Overall, these results help explain how the treatment with Abd-VHL has an effect on cell division in the LMO2 expressing cells.

**Figure 5.** Proteomic profiling of T-ALL cells after treatment with Abd PROTAC degrader. LMO2 expressing T cells (KOPT-K1) and LMO2 non-expressing T cells (Jurkat and RPMI8402) were treated in duplicate with 15 μM Abd-VHL for 24 hours. Panel **A**: Floating bar plots of LMO2 protein levels measured by targeted proteomics in control (DMSO treated) (grey bars) and Abd-VHL treated (red bars) in KOPT-K1, Jurkat and RPMI8402 cells. Panel **B**: Western blotting analysis of LMO2 levels in KOPT-K1 cells when treated with DMSO only and Abd-VHL for 24 hours. β-actin protein detection was used as an internal loading control for Western blotting analysis. Panel **C**: Heatmap displaying enriched terms from GSEA in KOPT-K1, Jurkat, and RPMI8402 cells after Abd-VHL treatment compared to control treatment (DMSO), based on global proteomics analysis. The enrichment score ranges from -2.5 (blue) to 2.5 (yellow). Panel **D** and **E:** Volcano plot of proteomic changes following Abd-VHL treatment compared to control treatment (DMSO) in KOPT-K1 (**D**) and Jurkat (**E**). The plots are based on the fold change (log_2_FC) and p-value (-log_10_(p-value)). Respectively, the yellow or grey circles indicate proteins with statistically significant or non-significant up-regulation/down-regulation. The cut-off for the volcano plots is log_2_FC=0.585, p-value=0.05). Panel **F**: a tabulation of statistically significant proteins in KOPT-KI, represented as yellow circles in Panel D compared to Jurkat cells. The data are presented as fold change (log_2_FC) and p-value (-log_10_(p-value)).

### Loss of the LMO2 protein complex initiates apoptosis

We investigated whether LMO2 protein degradation in the T-ALL cell lines resulted in programmed cell death (PCD) that accompanies growth inhibition effects. KOPT-K1 or Jurkat cells were treated with a single 20 μM dose of either Abd-CRBN or Abd-VHL for up to 30 hours and caspase 3/7 activation determined at the various time points. Increased levels of caspases were found in LMO2-expressing KOPT-K1 treated with either Abd compound, rising 10 fold (from 23325 RLU at 4 hours to 228865 RLU at 30 hours) using Abd-CRBN and 25 fold increase (from19460 RLU at 4 hours to 476843 RLU at 30 hours) using Abd-VHL (Fig. 6A). This rise in activated caspases mirrors loss of LMO2 as judged by Western blotting analysis (Fig. 6B) and with appearance of cleaved caspase 3, cleaved caspase 7 and cleaved poly(ADP-ribose) polymerase (PARP). The increase of caspase 3/7 activity in KOPT-K1 cells treated with Abd-CRBN is lower than observed with Abd-VHL, reflected by relatively low levels of protein detected in Western blot.

**Figure 6.**
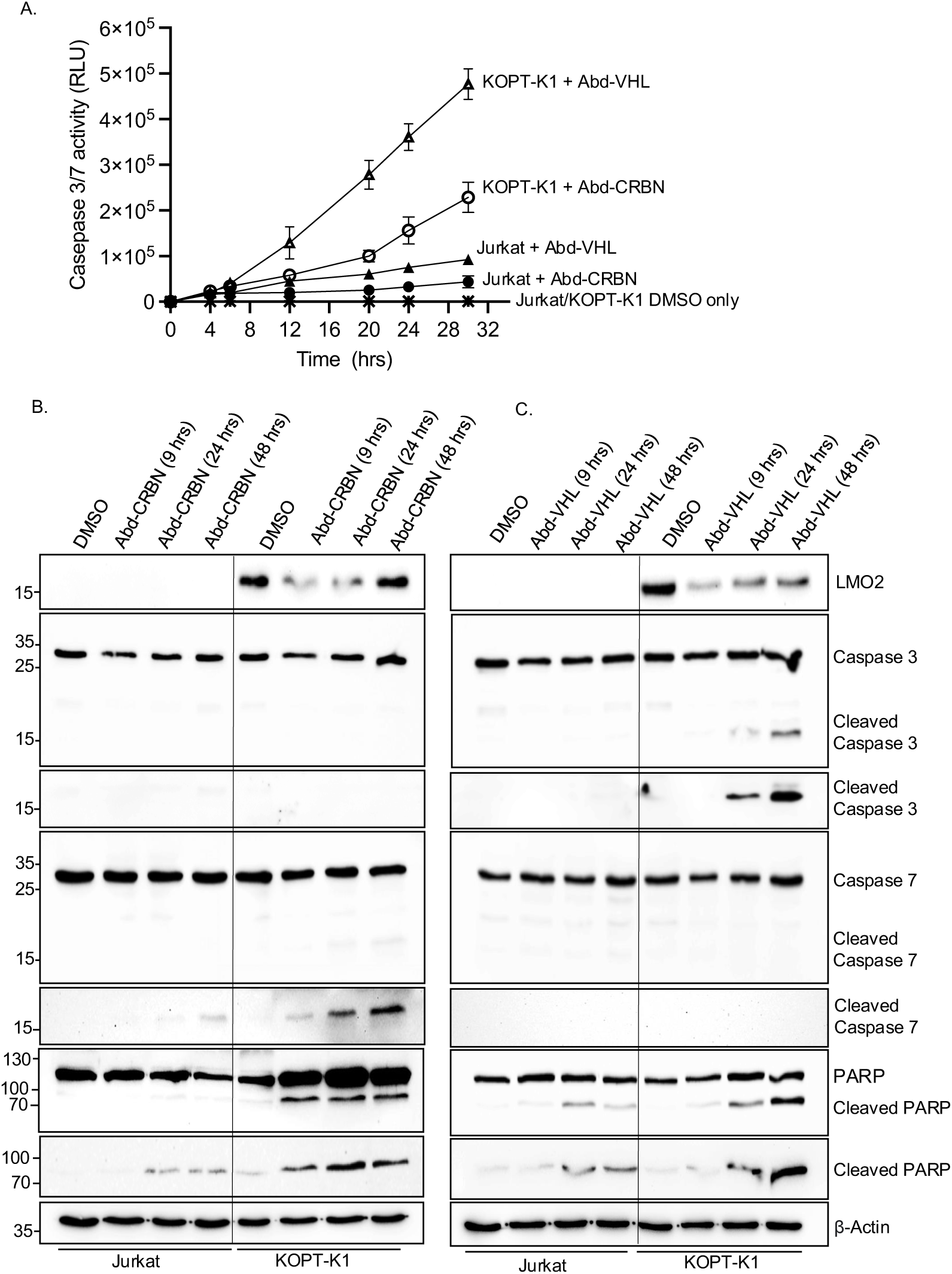
Caspase and PARP cleavage in cells after treatment with Abd degraders indicative of apoptosis initiation. KOPT-K1 and Jurkat cells were treated with Abd-CRBN or Abd-VHL and effects on viability and programmed cell death were analysed at different times. Panel **A:** Time course of progressive expression of caspases after the treatment with compounds was assayed for caspase 3/7 levels using Caspase-Glo^®^ 3/7. Cells were treated with 20 μM Abd-CRBN or Abd-VHL followed by cell culture up to 30 hours. Cells treated with 1% DMSO in culture medium was used as control. Data represent mean ± SEM (n=3). KOPT-K1 and Jurkat cells were treated with 20 μM Abd-CRBN (panel **B)** or Abd-VHL (panel **C)** and cultured for 24 or 48 hours. Cell extracts were made and proteins subjected to Western blotting analysis with specific antibodies for detection of LMO2, caspase 3, cleaved caspase 3, caspase 7, cleaved caspase 7, PARP, and cleaved PARP. β-actin protein detection was used as an internal loading control for Western blotting analysis.

Conversely, the effect of both compounds on the LMO2-negative Jurkat cells was a small increase in caspase 3/7 levels over the time course of the experiment. This increased 2.6 fold (from 16223 RLU at 4 hours to 43651 RLU at 30 hours) using Abd-CRBN and 4.5 fold (from 20309 RLU at 4 hours to 92683 RLU at 30 hours) using Abd-VHL. Western blotting analysis up to 48 hours of treatment with compounds did not show significant levels of activated, cleaved caspase 3 and only small amounts of activated, cleaved caspase 7 at 48 hours after treatment with Abd-CRBN (Fig. 6B). Similarly, Abd-VHL treatment (Fig. 6C) of Jurkat cells did not induce detectable cleaved caspase 3 or 7 in the Western blotting analysis. Small amounts of cleaved PARP were detected in the Western blotting analysis for treatment with both Abd compounds. Cells treated with only DMSO in culture medium showed essentially no detectable activated caspase 3/7 during this growth period (Fig. 6A) nor activated, cleaved caspase or PARP (Fig. 6B and C). To corroborate the detection system for cleaved caspase 7, both Jurkat and KOPT-K1 cells were treated with the chemotherapy reagent doxorubicin and analysed by Western blotting analysis (Supplementary Fig. 9A) and using both the luminescence assay (Supplementary Fig. 9B).

## DISCUSSION

### LMO2 depletion by antibody biodegraders or PROTACs interferes with T-ALL

Transcription factors like LMO2 have been considered as hard to drug targets because they are involved in protein-protein or protein-DNA interaction and often have large intrinsically disordered regions, as in the case of LMO2 (16). Therefore, using intracellular antibodies as inhibitors is a convenient starting point for drug discovery with these IDPs. In this study, we have shown that protein degradation of the LMO2 transcription factor occurs after either a binary interaction of an iDAb-E3 ligase fusion with LMO2 or a ternary interaction through PROTACs. Using an intracellular antibody for study of IDPs offers unique options because of the natural property of antibodies of high discriminating power, they can be easily manipulated by protein engineering (35) and can be used for target validation as a step before embarking on small molecule drug development campaigns.

The function of LMO2 to regulate cell proliferation has been found involved during tumorigenesis and the inhibition of LMO2 showed apoptosis induction in breast and colorectal cancer (36). In acute myeloid leukemia (AML) cells, the LMO2/LDB1 complex has an important role to promote cell growth and proliferation and is required for cells survival (37). Our data show that the depletion of LMO2 has an effect on growth of T-ALL expressing LMO2 but that the compounds have no off-target effect on non-expressing LMO2 T-ALL cells. Proteomic analysis (Fig. 5) exploring the pathways associated with LMO2 protein reduction in T-ALL cells supports the conclusion that LMO2 downregulation has an effect proteins required for cell division, coupled to down regulation of components of the ubiquitination machinery, known to coordinate phases and progression of cell division (38). Suppression of ubiquitination reduces proliferation and induces apoptosis, which was observed in treated cells (Fig. 6). Further, several proteins involved in mitosis are reduced upon LMO2 degradation (annotated in Supplementary Fig. 10), such CCNA2 (Cyclin A2), which controls G1/S and the G2/M transitions (38), is reduced in the LMO2 PROTAC treated cells. In relation to ubiquitination itself and protein turnover, it is significant that both UBE2C and UBE2S E2 ubiquitin enzymes are reduced in LMO2 depleted cells. These two E2 enzymes ubiquitinate anaphase-promoting complex/cyclosome (APC/C) required proteosomal loss of cyclin B1 for exiting from mitosis (39). Overall, the data suggest that when LMO2 is depleted from the T-ALL cells, the effect is to block mitosis and apoptosis follows.

### Collateral turnover of transcription factor partners with LMO2 proteosome degradation

The LMO2 protein is a transcription factor that operates as a bridging molecule for a multiprotein complex that binds to DNA through bipartite regions comprising a bHLH heterodimer of TAL1 and E47 (binding to E box elements) and a GATA protein (binding to GATA elements) (13). While LIM only proteins have two LIM domains, each with two LIM zinc binding fingers, these proteins do not interact directly with DNA. The LMO2 protein complex is required for function within T-ALL and forced expression of LMO2 and TAL1 in transgenic mouse models demonstrated this leukaemogenesis cooperativity (25). However, in the experimental system described here, we have demonstrated the effects on cell growth by treating T cells with the LMO2 PROTACs and both TAL1 and E47 were rapidly and simultaneously degraded with LMO2 (Figs. 1 and 3). Since there is a close association of the two bHLH with LMO2 proteins within the protein complex, lysines in these two proteins that lie close to the PROTAC binding site (as illustrated in Supplementary Fig. 11) could also undergo ubiquitination. Such an effect was described as bystander ubiquitination (40). Alternatively, degradation of LMO2 could destabilise the whole complex, in a natural process of transcription complex regulation, causing degradation of TAL1 and E47, as previously reported with PROTACs targeting the BAF chromatin remodeling complex (41). In addition, the half-lives of proteins in LMO2 protein complex were examined using cycloheximide to block protein synthesis. The result showed that TAL1 and E47 have a short half-life (10 minutes and 1 hour, respectively in this assay, Supplementary Fig. 12) and LMO2 protein has a half-life around 6 hours (42).

We could not find evidence of differential loss of other members of the complex (LDB1 and GATA) or LYL1 that is a different bHLH protein implicated in T-ALL. This may be because bystander ubiquitination could not occur due to the interaction surfaces of these proteins in relation to the PROTAC docking sites. Alternatively, the disparity in the degradation of LMO2, TAL1 and E47 compared with LDB1 and GATA proteins may be attributed to different pool sizes for LMO2-bound and free components of the transcription complex. The loss of TAL1 and E47 bHLH proteins along with LMO2 in the degradation cascade is, however, a complementary effect that enhances the effectiveness of anti-LMO2 PROTAC compounds.

### A strategy for drug discovery against intrinsically disordered proteins

Either of the two compounds described here may be developed for specifically treating T-ALL expressing LMO2. Furthermore, our data also illustrate a strategy for drug discovery for intrinsically disordered proteins, estimated that up to 30% cellular proteins have large intrinsically disordered regions (43), for instance transcription factors. Starting with a domain antibody that had been used for LMO2 target validation (15), compound surrogates were discovered using the antibody as a screening tool (17) and the compounds developed using medicinal chemistry. This is a general approach for hard to drug, disordered proteins, such as transcription factors.

## MATERIALS AND METHODS

Antibodies and reagent sources are found in Supplementary Table 1.

### Molecular cloning

The iDAb LMO2-E3 ligases fusions were generated by polymerase chain reaction (PCR) using Phusion^®^ High-Fidelity PCR kit (NEB). pEF-VH576-L10-FLAG-CRBN-mb was used as a template to generate VH576-L10-FLAG-CRBN; pEF-FLAG-CRBN-L5-VH576-myc was used as a template to generate FLAG-CRBN-L5-VH576; pEF-VHL-L5-FLAG-VH576-mb was used as a template to generate VHL-L5-FLAG-VH576; pEF-NtFarn-FLAG-VH576-L10-VHL was used as a template to generate FLAG-VH576-L10-VHL; pEF-UBOX-L5-FLAG-VH576-mb was used as a template to generate UBOX-L5-FLAG-VH576; pEF-NtFarn-FLAG-VH576-L5-UBOX was used as a template to generate FLAG-VH576-L5-UBOX. All the PCR templates were previously described (17). PCR products were cloned into pEF-BOS plasmid and sequenced to confirm inset with the vector.

Generation of LMO2, MAX, RHOBWT-FL, KRASWT-FL, GFP^2^, ISL-1, and Zn268 was by PCR using Phusion^®^ High-Fidelity PCR kit (NEB) and cloned into pEF-BOS-Rluc8 plasmid to produce pEF-LMO2-Rluc8, pEF-MAX-Rluc8, pEF-Rluc8-RHOBWT-FL, pEF-Rluc8-KRASWT-FL, pEF-GFP^2^-Rluc8, pEF-ISL-1-Rluc8, and pEF-Zn268-Rluc8 (17, 44).

### Cell culture

Human T-ALL cell lines KOPT-K1, Jurkat, MOLT-3, ALL-SIL, DND-41, PF382, SUPT-1, RPMI8402, and P12-Ichikawa cell lines were cultured in RPMI-1640 medium (Gibco^TM^) supplemented with 10% foetal bovine serum (FBS) (Pan-Biotech). HEK293T cell line was cultured in DMEM (Gibco^TM^) supplemented with 10% FBS. Cells were grown at 37°C in a humidified incubator with 5% CO_2_ and were regularly performed mycoplasma test using a MycoAlert Mycoplasma Detection kit (Lonza). Cells with mycoplasma-free were used for experiments.

### Transient Transfection

HEK293T cells were seeded on 6-well plate at 3x10^5^ cells/well and incubated at 37°C with 5% CO_2_ until 70-80% confluence determined visually. After 24 hours, cells were transfected with Lipofectamine2000 reagent (Invitrogen) according to manufacturer’s instructions. An iDAb LMO2-E3 ligase plasmid (ranging from 0-1.0 μg) was mixed with 1 μg pEF-BOS-LMO2 expression vector (45) and diluted in 150 ul Opti-MEM reduced serum medium (Gibco^TM^). 10 μl Lipofectamine2000 was diluted in 150 ul Opti-MEM^TM^ reduced serum medium. The diluted DNA was added to diluted Lipofectamine2000 and incubated for 15 minutes at room temperature. After the incubation, the mixture was added to the HEK293T cells. After 24 hours of transfection, the transfected cells were detached from the plate using trypsin-EDTA (0.05%), phenol red (Gibco^TM^) and incubated at 37°C for 2 minutes. Two volumes of pre-warmed complete media were added to inactivate trypsin, and dispersal of the medium, by pipetting over the cell layer surface, was used to recovery of >95% of cells. Cell pellets were collected by centrifugation at 100g for 5 minutes at room temperature.

### Luciferase reporter assay

HEK293T cells were transiently transfected with 500 ng reporter vectors (pEF-LMO2-Rluc8, pEF-MAX-Rluc8, pEF-RHOBWT-FL-Rluc8, pEF-KRASWT-FL-Rluc8, pEF-GFP^2^-Rluc8, pEF-ISL-1-Rluc8, or pEF-Zn268-Rluc8). After 24 hours, the transfected cells were re-seeded in white opaque 96-well plate at 5x10^4^ cells/well and incubated at 37°C for 4 hours prior the compound addition. The treated cells were incubated at 37°C for 24 hours, after which 10 μl of 250 μM luciferase substrate (Coelenterazine 400a) (Cayman Chemical) were added per well to get 10 μM at final concentration. The luminescence signal was measure at 400-700 nm wavelength filter luminescence using PHERAstar^®^ FSX (BMG Labtech).

### Lentiviral cell lines

The lentiviral transfer vector plasmid TLCV2-VH576-L10-FLAG-CRBN was constructed with TLCV2 (26). The pMA-VH576-L10-FLAG-CRBN was synthesised from GeneArt (ThermoFisher). The insert was removed from the pMA vector by digesting with AgeI and NheI, then cloned into TLCV2 vector between Age/NheI sites to produce a TLCV2-VH576-L10-FLAG-CRBN plasmid.

Lentiviral particles were produced by transient co-transfection of lentiviral transfer vector plasmid and the packaging plasmids (pRSV-Rev, pMDLg/pRRE, and pMD2-VSV-G) in HEK293T cells using Lipofectamine2000. After 48 hours transfection, debris was removed by spinning at 300g for 5 minutes at 4 °C, followed by filtration of the supernatants through 0.45 μm filter to remove viral aggregates. The filtered lentiviral supernatants were centrifuged at 15,000g for 24 hours at 4 °C. Virus pellets were resuspended in completed media and stored at -80 °C in cryotubes. The lentiviral particles were titrated by serial dilution (3x dilution) in complete media. After 48 hours post-transduction, cells were harvested and analysed by flow cytometry for the viral GFP reporter expression and the viral titres calculated.

### T cell transduction

KOPT-K1 and Jurkat cells were infected by using TransDux MAX^TM^ virus transduction reagent and followed the spinoculation method (SBI, System Bioscience). For the spinoculation, KOPT-K1 and Jurkat were seeded in 24-well plate at 5x10^5^ cells/well and resuspended with 400 μl complete RPMI media plus 100 μl MAX Enhancer, 2 μl TransDux^TM^, and 4 μl 1M HEPES pH7.0 buffer. Virus suspensions were added to cells at a multiplicity of infection of 3 and the plate was centrifuged at 1,500g at 32 °C for 2 hours. After spinoculation, the presence of the cells was verified at the bottom of the wells and 400 μl complete medium was added to each well. Cells resuspended by pipetting. The cell suspensions were transferred to 1.5 ml sterile tubes and centrifuged at 1,500g for 5 mins at room temperature. The supernatant was discarded to remove the transduction reagent, cells resuspended in 400 μl fresh complete media and transferred to a new 24-well plate. The infected cells were incubated at 37 °C, 5% CO_2_ incubator for 48 hours. The doxycycline was added to induce the biodegrader expression before analysis by flow cytometry and Western blotting.

### Western blotting

Cells were washed with PBS before lysis with RIPA buffer (Sigma-Aldrich) (50mM Tris-HCL, pH 8.0, 150mM sodium chloride, 1.0% Igepal CA-630 (NP-40), 0.5% sodium deoxycholate (SDC), and 0.1% sodium dodecyl sulphate) supplemented with Pierce^TM^ protease inhibitor tablets, EDTA-free (ThermoFisher). Cells were lysed on ice and vortexed every 5 minutes for 30 mins, then centrifuged at 10,000g, 10 minutes at 4 °C. Protein concentrations were quantified using Pierce^TM^ BCA Protein assay kit (ThermoFisher) with the BCA standard curve range from 0-2 mg/ml. Equal amounts of protein samples were separated on 10% or 15% SDS-PAGE and subsequently transferred to polyvinylidene fluoride (PVDF) membrane (Amersham, Cytiva) by wet blotting method. The membrane was blocked with 10% non-fat milk (Sigma-Aldrich) in PBST (PBS with 0.1% Tween20 (Sigma-Aldrich)) before incubation with primary antibodies overnight at 4°C on a roller. After the incubation, the membrane was washed with PBST for 5 minutes and repeating washing step for 5 times. The washed membrane was incubated with horseradish peroxidase-conjugated secondary antibodies at room temperature for 1 hour on a roller and washed with PBST for 5 minutes and repeating the washing step for 5 times. The membrane was incubated with Clarity^TM^ western blotting substrate (Bio-Rad) for 1 minute before exposure to a ChemiDoc imager (Bio-Rad). For quantification, densitometry analysis of protein expression was analyzed using Image Lab software.

### Reverse transcription-PCR (RT-PCR)

The expression of *LMO1* and *LMO2* was verified using RT-PCR method using the primers shown in Supplementary Fig. 8. 1x10^7^ cells were harvested and lysed by using 0.5% NP40 lysis buffer (Thermo Scientific). Total cellular RNA was extracted from the supernatant by using water-saturated phenol pH6.6 (Invitrogen^TM^), 2M NaAc pH5.0 was added to 300mM and 100% ethanol to 70% to precipitate RNA at -20°C for 2 hours. The pellet was washed with 70% aqueous ethanol, dried at room temperature. The RNA precipitate was dissolved in 10mM Tris pH7.5. The final RNA concentration was measured using Nanodrop^TM^ spectrophotometer (Thermo Scientific). Two μg of RNA was used for first strand cDNA synthesis with 100 pmoles oligo(dT)_12-18_ primer (Invitrogen^TM^) and 20 units SuperScript^TM^ II Reverse Transcriptase (Invitrogen^TM^). Primers for LMO1 were forward 5’-GATCCAGCCCAAAGGGAAGCAG-3’, reverse 5’-GATAAAGGTGCCATTGAGCTG-3’.

Primers for LMO2 were forward 5’-GATTCCTCGGCCATCGAAAGG-3’, reverse 5’-GATGTTTGTAGTAGAGGCGCCG-3’. Primers for KRAS were forward 5’-GATATGAC TGAATATAAACTTGTGGTAG-3’, reverse 5’-GATGGCAAATACACAAAGAAAGC-3’.

PCR was performed by using DNA Engine Tetrad^®^ Thermal cycler (Bio-Rad). The PCR conditions were 98°C 1 minute 1 cycle for initial denaturation, and 30 cycles of 98°C 10 seconds for denaturation, 65°C 15 seconds for annealing, 72°C 30 seconds per 1 kb for extension, then 72°C 10 minutes for final extension. The PCR products were resolved on 1.0% agarose gel, visualized by SYBR safe staining, and quantified using ChemiDoc imager (Bio-Rad).

### Genomic PCR

The presence of the chromosomal translocation t(11;14)(p13;q11) in KOPT-K1 cells (27) was confirmed using genomic PCR. The derived genomic translocation sequence is shown in Supplementary Fig. 13. Total DNA was extracted from 1x10^6^ KOPT-K1 cells by using DNeasy Blood&Tissue kit (Qiagen) according to the manufacturer’s instruction. The final DNA concentration was measured using Nanodrop^TM^ spectrophotometer (Thermo Scientific). One hundred ng of DNA template was used to generate PCR using Phusion^®^ High-Fidelity PCR kit (NEB). PCR was performed by using DNA Engine Tetrad^®^ Thermal cycler (Bio-Rad). The forward and reverse primers used to confirm the chromosomal translocation in KOPT-K1 was forward two 5’-GATGAAT TCGAAGCTACTGCAGCCATC-3’, forward three 5’-GATGAATTCATGCTATGAGGTA GGTATG-3’, and J delta reverse 5’-GATGGATCCGGTTCCACAGTCACTCG GGTTCC-3’. The PCR condition was 98°C 1 minute 1 cycle for initial denaturation, followed by 98°C 10 seconds for denaturation, 65°C 15 seconds for annealing, 72°C 15 seconds for extension for 30 cycles, then 72°C 10 minutes for final extension. The PCR products were resolved on 1% agarose gel, visualized by SYBR safe staining, and quantified using ChemiDoc imager (Bio-Rad).

### Chemistry; synthesis, purification, and quality control

Details appear in the Supplementary information file.

### Cell assays with Abd-CRBN, Abd-VHL and inhibitors

Assays for the LMO2 PROTAC compounds Abd-CRBN and Abd-VHL were carried out with compounds synthesized by contract with O2H Discovery. All compounds (Abd compounds and epoxomicin, thalidomide, VHL, and MLN4924) were dissolved in 100% DMSO at 10mM and stored in small aliquots -20°C. The stability of the compounds was evaluated before using for experiments by using Liquid Chromatography-Mass Spectrometry (LC-MS) method using a Xevo TQ Mass spectrometer instrument.

### Cell viability assays

The number of viable cells was measured by staining cells with 0.4% Trypan blue solution (Invitrogen^TM^) and counting with Countess^TM^ Automated Cell Counter (Invitrogen^TM^). For inhibition studies, KOPT-K1 cells were seeded in 6-well plate at 5x10^5^ cells/well. After the incubation at 37°C in a humidified incubator with 5% CO_2_ for 24 hours, cells were pre-treated with inhibitor compounds (epoxomicin, thalidomide, or lenalidomide) or Neddylation inhibitor (MLN4924) for 2 hours or proteasome inhibitor (epoxomicin) for 24 hours prior the treatment of Abd compounds for 24 hours. For apoptosis studies, KOPT-K1 cells were seeded in 6-well plate at 5x10^5^ cells/well. After the incubation at 37°C in a humidified incubator with 5% CO_2_ for 24 hours, cells were treated with Abd-CRBN or Abd-VHL for 24 or 48 hours.

Cell viability was also assessed by CellTiter-Glo^®^ (Promega) according to the manufacturer’s instructions. Cells were seeded in triplicate in white 96-well plate (PerkinElmer) at 1x10^4^ cells/well. After the treatment with compounds, an equal amount of CellTiter-Glo^®^ reagent was added to each well and incubated at room temperature for 10 minutes before the luminescence signal measurement using PHERAstar^®^ FSX (BMG Labtech).

### Dose response and EC_50_ determination

KOPT-K1, Jurkat, and MOLT-3 cells were seeded in white 96-well plate at 1x10^4^ cells/well and incubated at 37°C in a humidified incubator with 5% CO_2_. After 24 hours incubation, cells were treated with 5, 10, 15, 20, or 25 μM of Abd-CRBN or Abd-VHL for 24 hours. The viable cells were measured using CellTiter-Glo^®^ assay. The EC_50_ was calculated using Prism9 GraphPad Software. For dose response by western blotting analysis, KOPT-K1 and Jurkat cells were seeded in 6-well plate at 5x10^5^ cells/well and the incubation at 37°C in a humidified incubator with 5% CO_2_ for 24 hours. Then, cells were treated with Abd-CRBN or Abd-VHL compounds at concentration of 5, 10, or 20 µM for 2, 6, or 24 hours.

### Caspase3/7 activity assay

Caspase activities were measured using Caspase-Glo^®^ 3/7 (Promega) according to the manufacturer’s instruction. Cells were seeded in triplicate in white 96-well plate (PerkinElmer) at 1x10^4^ cells/well. After the treatment with Abd-CRBN or Abd-VHL compounds for 24 hours, an equal amount of CellTiter-Glo^®^ reagent was added to each well and mixed using a plate shaker at 300 rpm for 30 seconds, incubated the plate at room temperature for 1 hour before the luminescence signal measurement using PHERAstar^®^ FSX (BMG Labtech).

### Proteomic Analysis by Mass Spectrometry

Samples were prepared using the SimPLIT workflow previously described (46) with minor modifications. Cell pellets were lysed in a buffer containing 0.1 M TEAB, 1% SDC, 10% isopropanol, 50 mM NaCl, 5 mM TCEP, 10 mM IAA, nuclease, and protease/phosphatase inhibitors. After 5 min of bath sonication and 45 min at room temperature, protein concentration was measured by Bradford assay. For digestion, 15 µg protein aliquots were treated with trypsin and incubated at 37 °C for 2 hours, then overnight at room temperature. Digested samples were acidified, spun, desalted, and labelled with TMTpro-18plex. Labelled samples were combined, acidified, centrifuged to remove SDC, and dried.

Offline peptide fractionation was based on high pH Reverse Phase (RP) chromatography using the Waters XBridge C18 column (2.1 × 150 mm, 3.5 μm) on a Dionex Ultimate 3000 HPLC as previously described (47) with minor modifications. Retention time-based fractions are collected and pooled into 24-samples for LC-MS analysis.

Samples were analyzed on a Dionex UltiMate 3000 UHPLC with an Orbitrap Ascend Tribrid mass spectrometer and a PepMap C18 capillary column, as previously described (47). A 120-minute gradient separation was applied. Data acquisition used TMT-SPS-MS3 with real-time database search, targeting precursors between 400–1600 m/z, and employing RTS spectral identification of MS2 fragments to trigger SPS-MS3 scans.

Targeted proteomics analysis by mass spectrometry was further conducted on the fractions containing LMO1 and LMO2 peptides detected in the global proteomics analysis to improve sensitivity. Precursors (593.01, 582.34, 693.68, 745.72, 888.52, and 921.39 m/z) were selected in the quadrupole with an isolation width of 0.7 m/z and fragmented with HCD using 36% collision energy (CE). MS2 spectra were recorded in profile mode in Orbitrap at 45,000 resolutions using a 20-scan mode for the isolation list and an AGC setting of 5 x 10^5^.

Mass spectra were analyzed using SEQUEST-HT and COMET in Proteome Discoverer 3.0 for protein identification and quantification. Parameters included a 20-ppm precursor mass tolerance, 0.5 Da for TMT-MS3, and 0.02 Da for targeted-MS2. Searches targeted fully tryptic peptides with up to 2 mis-cleavages, with TMTpro and carbamidomethylation as static modifications, and methionine oxidation and asparagine/glutamine deamidation as variable. Peptide confidence was estimated with Percolator, FDR set at 0.01 and validated by q value and decoy database search.

Proteomics data normalization, analysis, and visualization: Each TMT column was normalised by dividing the protein abundance values by the median of each value per column. The normalised data were used to calculate log2 ratios of treated vs DMSO for all cell lines. Gene Set Enrichment Analysis (GSEA) (48) was conducted to evaluate the biological effects of 2294 treatment by analysing log2 ratios of protein abundance from global proteomics data. The proteins were ranked based on log2 ratios, indicating upregulation or downregulation in 2294-treated samples compared to DMSO controls. Significant GSEA enrichment scores were calculated using the annotation terms using KEGG and GO annotation terms, applying a p-value threshold of < 0.05 with Benjamini-Hochberg adjustment. Significant terms were visualised using ridge plots and a heatmap summarising the top 10 terms based on the adjusted p-value.

## Acknowledgements

This work was financed by grants from Blood Cancer UK (BCUK 12051 and 19013), CR- UK (Project Development Award) and by ICR Core. We wish to thank Prof. Marc Mansour for providing for providing T-ALL cell lines (P12-Ichikawa, MOLT-4, PF-382, DND-41, and ALL-SIL) and very grateful to Teresa Palomero, Foon Wu-Baer, and Richard Baer for LOUCY, SUP-TI and RPMI8402 cells. We also thank various ICR colleagues for generously providing reagents, the ICR Flow Cytometry Facility for their support, services, and expertise and Dr. John Caldwell for advice on chemical synthesis and O2H Group for batch synthesis of the Abd degrader compounds.

## Author contributions

Originator of project: THR Funding acquisition: THR

Participated in research design: NS, CB, NB, AR, AM, THR.

Conducted experiments: NS, CB, NB, AM.

Performed data analysis: All authors.

Contributed to the writing of the manuscript: All authors.

## Additional information

Supplementary Information accompanies this paper.

