## Supplementary table and figures for "Degradation of LMO2 in T cell leukaemia results in collateral breakdown of transcription complex partners and causes LMO2-dependent apoptosis"

### SUPPLEMENTARY INFORMATION

Supplementary Table
Supplementary Figures
Supplementary Information Chemical Synthesis

**SUPPLEMENTARY TABLE**

| Reagent type | Designation | Source | Identifiers |
|---|---|---|---|
| Abd degrader/PROTAC | Abd-VHL | o2h group | N/A |
| Abd degrader/PROTAC | Abd-CRBN | o2h group | N/A |
| Antibody | LMO2 goat antibody | R&D systems | Cat#AF2726 |
| Antibody | pan-RAS mouse antibody | Sigma-Aldrich | Cat#OP40 |
| Antibody | CRBN rabbit antibody | Cell Signaling | Cat#71810 |
| Antibody | VHL rabbit antibody | Invitrogen | Cat#PA5-27322 |
| Antibody | PARP rabbit antibody | Cell Signaling | Cat#9542 |
| Antibody | Caspase-3 rabbit antibody | Cell Signaling | Cat#9665 |
| Antibody | TAL1 rabbit antibody | A gift from Prof. Richard Bear (Columbia University) | N/A |
| Antibody | E2A (E47) rabbit antibody | Cell Signaling | Cat#4865 |
| Antibody | GATA-3 rabbit antibody | Cell Signaling | Cat#5852 |
| Antibody | LLY1 rabbit antibody | Invitrogen | Cat#PA5-68772 |
| Antibody | LDB1 rabbit antibody | Abcam | Cat#96799 |
| Antibody | Cleaved Caspase-3 rabbit antibody | Cell Signaling | Cat#9664 |
| Antibody | Caspase-7 rabbit antibody | Cell Signaling | Cat#9492 |
| Antibody | Cleaved Caspase-7 rabbit antibody | Cell Signaling | Cat#9491 |
| Antibody | FLAG M2 mouse antibody | Sigma-Aldrich | Cat#F1804 |
| Antibody | β-Actin mouse antibody | Sigma-Aldrich | Cat#A5441 |
| Antibody | Cyclophilin-B rabbit antibody | Abcam | Cat#ab178397 |
| Antibody | α-Tubulin rabbit antibody | Abcam | Cat#ab4074 |
| Antibody | Anti-Goat IgG HRP-linked antibody | Abcam | Cat#ab6741 |
| Antibody | Anti-Rabbit IgG HRP-linked antibody | Cell Signaling | Cat#7074S |
| Antibody | Anti-Mouse IgG HRP-linked antibody | Cell Signaling | Cat#7076S |
| Chemical compound | von Hippel-Lindau (VHL) Ligand 1 | Cayman chemical | Cat#21591 |
| Chemical compound | Cereblon (CRBN) | A gift from Habib Bouguenina | N/A |
| Chemical compound | Thalidomide | Sigma-Aldrich | Cat#T144 |
| Chemical compound | Lenalidomide | A gift from Habib Bouguenina | N/A |
| Chemical compound | Epoxomicin | A gift from Habib. Bouguenina | N/A |
| Chemical compound | MLN4924 | R&D systems | Cat#6499 |
| Chemical compound | Doxorubicin | Sigma | Cat#D9891 |
| Reagent kit | CellTiter-Glo® | Promega | Cat#G8090 |
| Reagent kit | Caspase-Glo®3/7 | Promega | Cat#G7570 |
| Reagent kit | Pierce™ BCA Protein assay kit | Thermo Scientific | Cat#23225 |

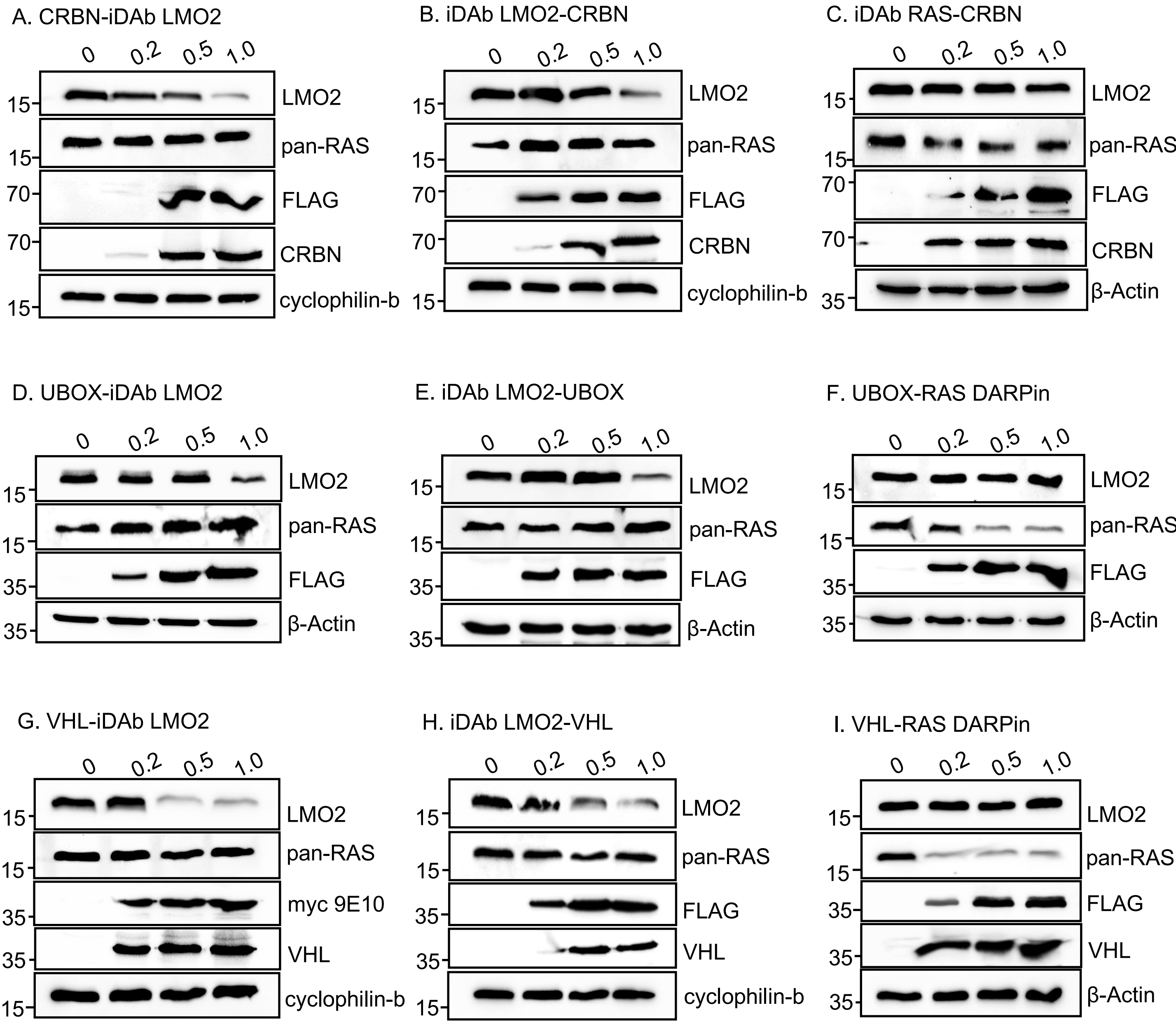


**Figure S1: LMO2 protein degradation in HEK293T cells after transfection with biodegrader constructs**

HEK293T cells were co-transfected with 1 µg pEF-BOS-LMO2 and plasmids expressing different iDAb LMO2-E3 ligase fusions for 24 hours. The amount of the iDAb LMO2-E3 ligase plasmids was titrated from 0 to 1.0 µg. pEF-BOS (vector plasmid) was used to equalise the amount of plasmid transfection to 2 ug. Western blotting analysis showing the expression of LMO2 in cells transfected with CRBN-iDAb LMO2 (panel A), iDAb LMO2-CRBN (panel B), anti-RAS iDAb-CRBN (panel C), UBOX-iDAb LMO2 (panel D), iDAb LMO2-UBOX (panel E), UBOX-anti-RAS DARPin (panel F), VHL-iDAb LMO2 (panel G), iDAb LMO2-VHL (panel H), and VHL-anti-RAS DARPin) (panel I). The antibodies used for protein detection are shown to the right of each panel. The anti-FLAG and anti-myc 9E10 antibodies detect the FLAG and myc tag on each iDAb-E3 ligase, the anti-CRBN antibody detects the transfected iDAb-CRBN in panels A-C and anti-VHL detects the transfected iDAb-VHL in panels G-I. Loading control proteins were detected with either anti-cyclophilin-b or anti-β-actin.

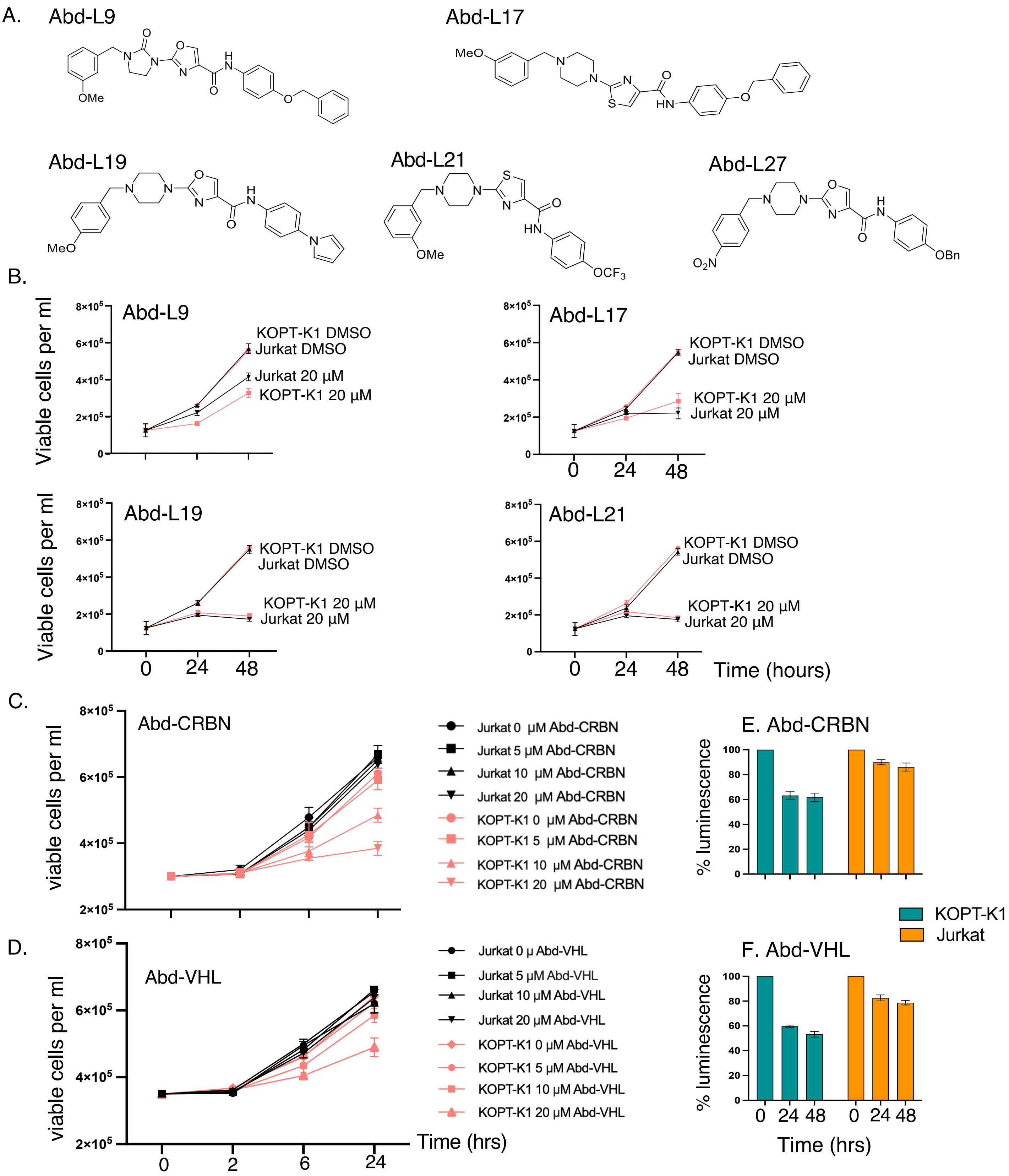


**Figure S2: Viability of Jurkat and KOPT-K1 cells after treatment with Abd compounds and Abd degraders**

Panel A: Chemical structures of LMO2 Abd compounds. Panel B: Jurkat and KOPT-K1 cells were treated at 20 µM of Abd-compounds from 0 to 48 hours. Cell viability after the treatment was measured by counting viable cells after staining cells with trypan blue. Panel C and D: Cell viability was also determined using trypan blue after the treatment with different concentrations of Abd degrader PROTAC compounds (0, 5, 10, and 20 µM) for 2, 6, and 24 hours. Panel E and F: CellTiter-Glo viability assays of Jurkat and KOPT-K1 cells treated with a single 20 µM dose of Abd-CRBN (panel E) or Abd-VHL (panel F) for up to 48 hours. All the values are presented as the average viable cells per ml. Data represent mean ± SEM (n=3).

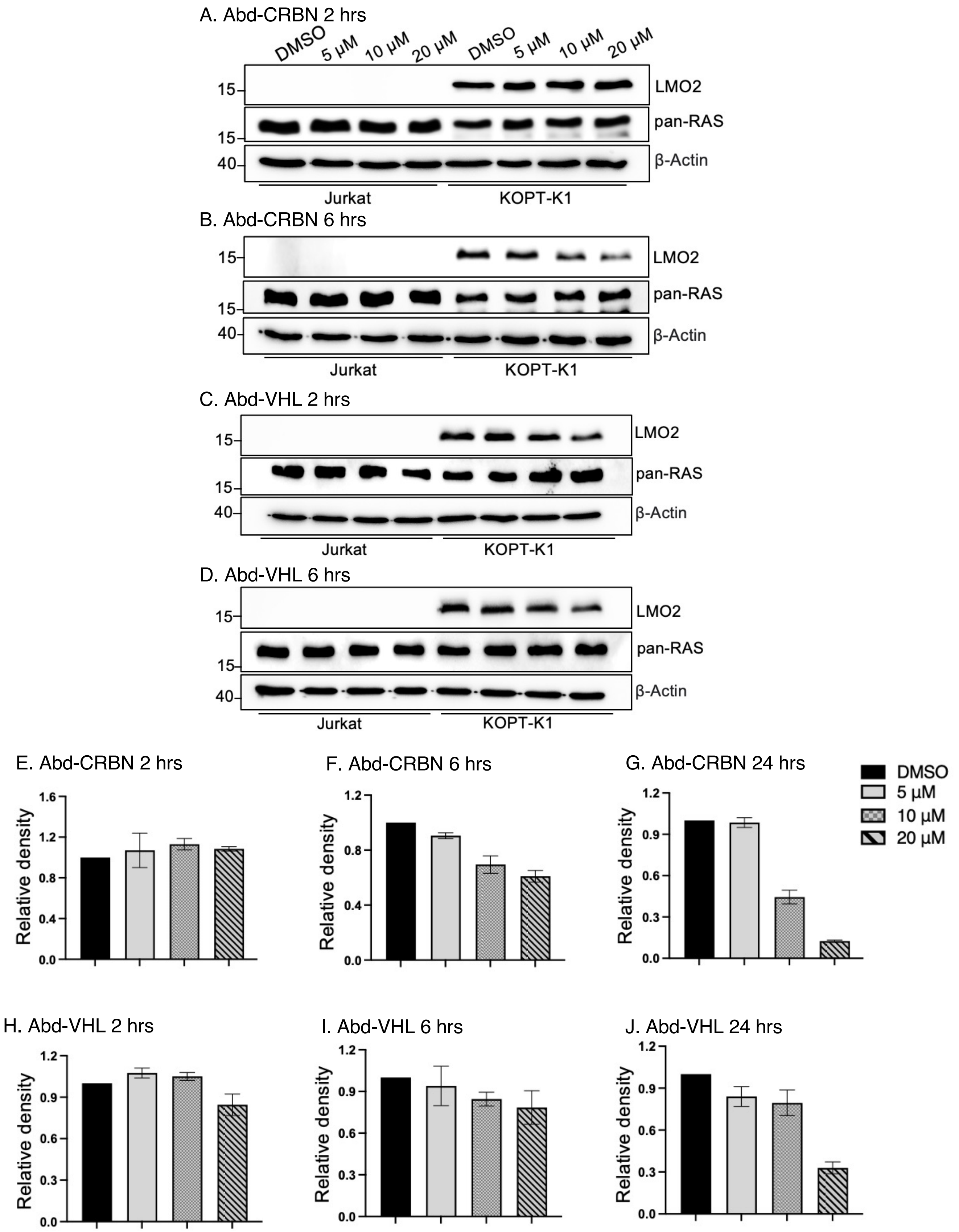


**Figure S3: LMO2 PROTEIN IN KOPT-K1 AND JURKAT CELLS AFTER TREATMENT WITH ABD COMPOUNDS**

KOPT-K1 and Jurkat cells were treated with different concentrations of Abd-degraders (at 0, 5, 10, and 20 μM) for 2, 6, and 24 hours and LMO2 protein levels determined by Western blot analysis. LMO2 protein in cells treated for 2 or 6 hours with Abd-CRBN (A, B), and Abd-VHL (C, D) or 24 hours (shown in Fig. 2, D, E). β-actin was used as an internal loading control for Western blotting analysis. Densitometry data of LMO2 expression in KOPT-K1 treated with Abd-CRBN (E,F,G) ir with Abd-CRBN (H, I, J) are presented as mean relative densitometry units. Data represent mean + SEM (n=3). $DC_{50}$ values of Abd-CRBN and Abd-VHL at 20 μM for 24 hours treatment in KOPT-K1 is 9 and 15 μM, respectively. The $DC_{50}$ was calculated using Graph Pad Prism 9.0 software.

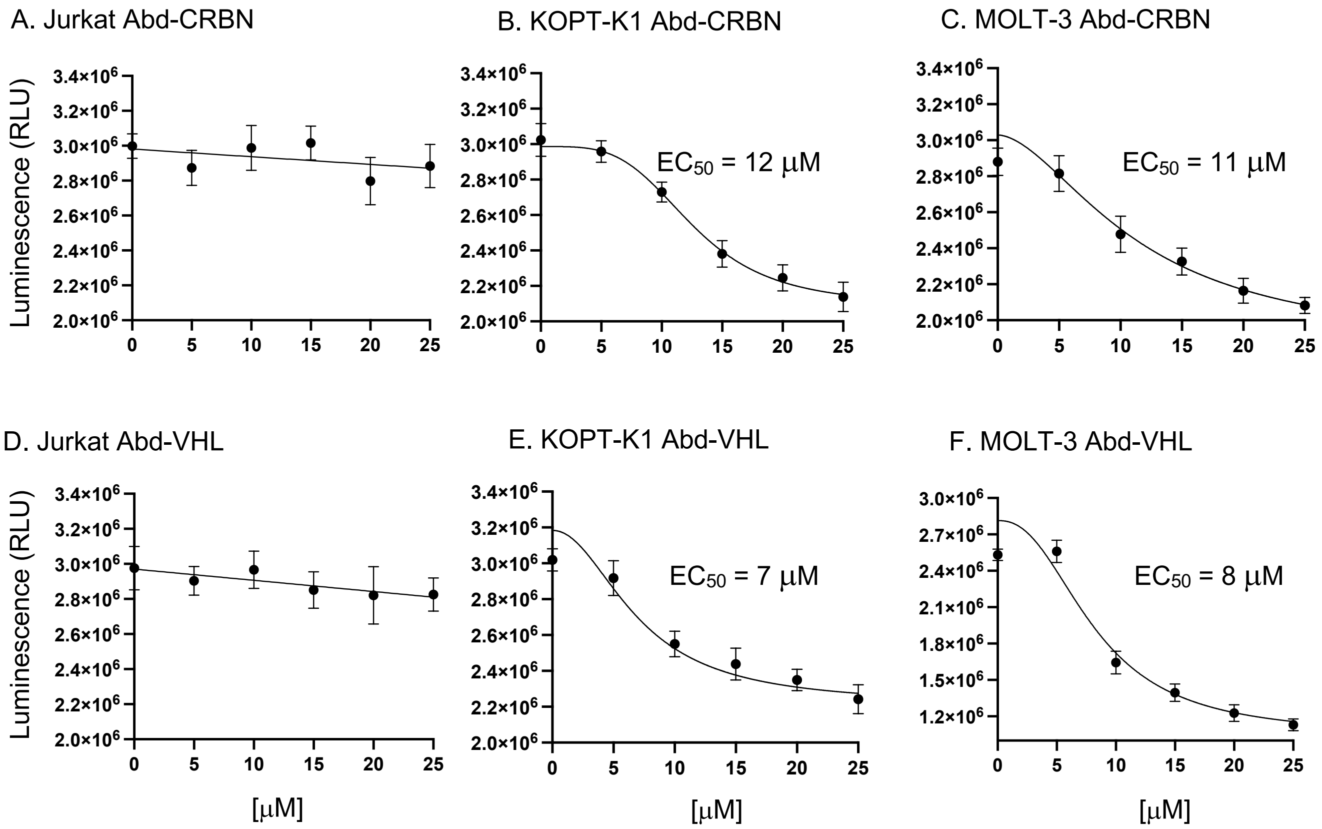


**Figure S4: Dose response of T cell lines with Abd degraders**

T cells were treated with Abd-CRBN or Abd-VHL in a dose range from 0-25 μM for 24 hours. The effect on viability was measured using CellTiter-Glo assays at 24 hours in Jurkat, KOPT-K1, and MOLT-3 with Abd-CRBN (panels A-C) or with Abd-VHL (panels D-F). Data represent mean ± SEM (n=3). $EC_{50}$ values were determined from dose-response curves generated using Graph Pad Prism 9.0 software.

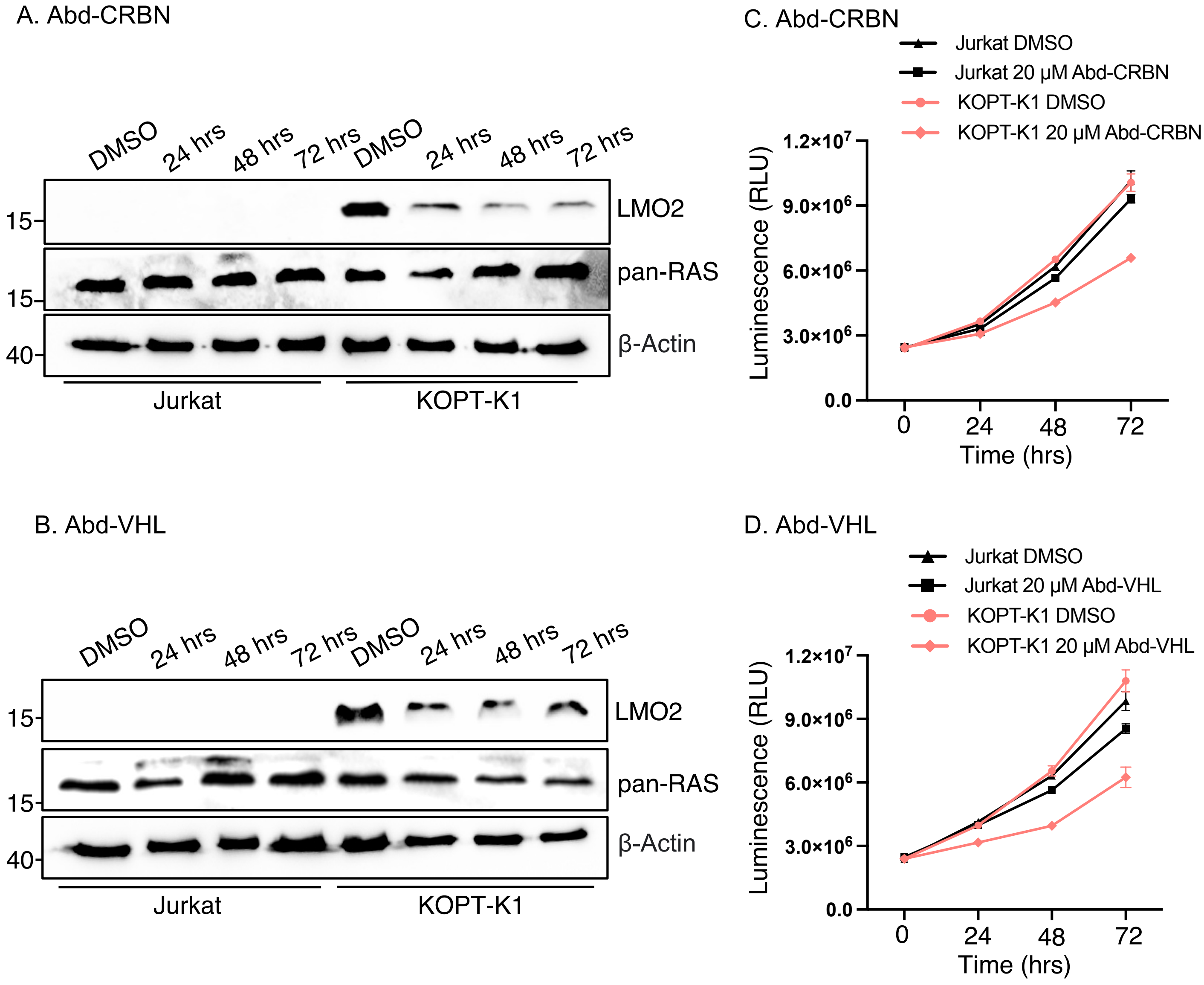


**Figure S5: Longevity of responses of KOPT-K1 cells to single dose treatment with Abd degraders** Jurkat and KOPT-K1 cells were treated with 20 µM of Abd-compounds and cultured from 0 to 72 hours. Western blot data of LMO2 and RAS expression in cells were treated with Abd-CRBN (panel A) and Abd-VHL (panel B). β-actin was used as an internal loading control for Western blotting analysis. Cell viability was determined using CellTiter-Glo assay after the treatment with Abd-CRBN (panel C) and Abd-VHL (panel D) at points during the 72 hours. All the values are presented as the average luminescence values. Data represent mean ± SEM (n=3).

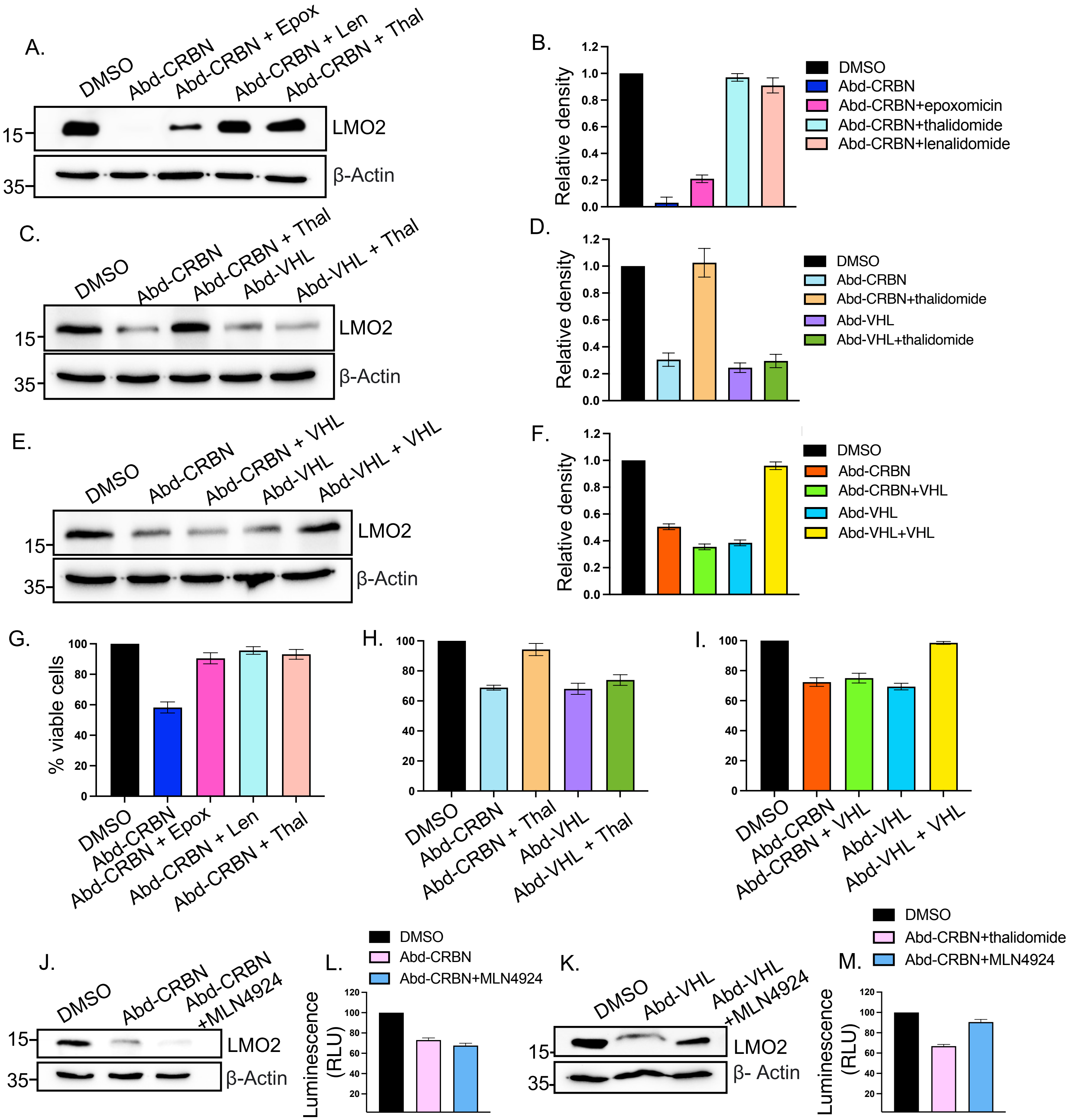


**Figure S6: LMO2 levels and viability of KOPT-K1 cells treated with Abd degrader compounds and E3 ligase inhibitors**

The involvement of proteasome machinery was investigated in KOPT-K1 with proteosome inhibitors or by competing with free E3 ligase ligands. Western blot data show LMO2 expression in KOPT-K1 cells after the treatment with or without inhibitors followed by Abd-compounds (panels A, C, E). Inhibitors used were the proteasome inhibitor epoxomicin (Epox) (0.8 µM), or CRBN inhibitors (10 µM thalidomide (thal), and 10 µM lenalidomide (len) (panels A, C) or free VHL ligand (panel E). Inhibitors added for 2 hours prior the treatment of 20 µM Abd-CRBN or Abd-VHL and further culture for 24 hours. β-actin was used as a loading control for analysis. Densitometry data of LMO2 in KOPT-K1 are presented as mean relative densitometry units, with standard deviation (panels B, D, F). After the treatment, cell viability was measured with trypan blue (panels G, H and I). Data represent mean ± SEM (n=3). KOPT-K1 cells were also treated with or without protein NEDDYlation inhibitor (10 µM MLN4941) for 2 hours prior the treatment of Abd-CRBN (panel J) or Abd-VHL (panel K) for 24 hours. CellTiter-Glo assays (panels L and M) was used to measure viability, presented as relative to luminescence in DMSO treated cells and normalized to 100%. Data represent mean ± SEM (n=3).

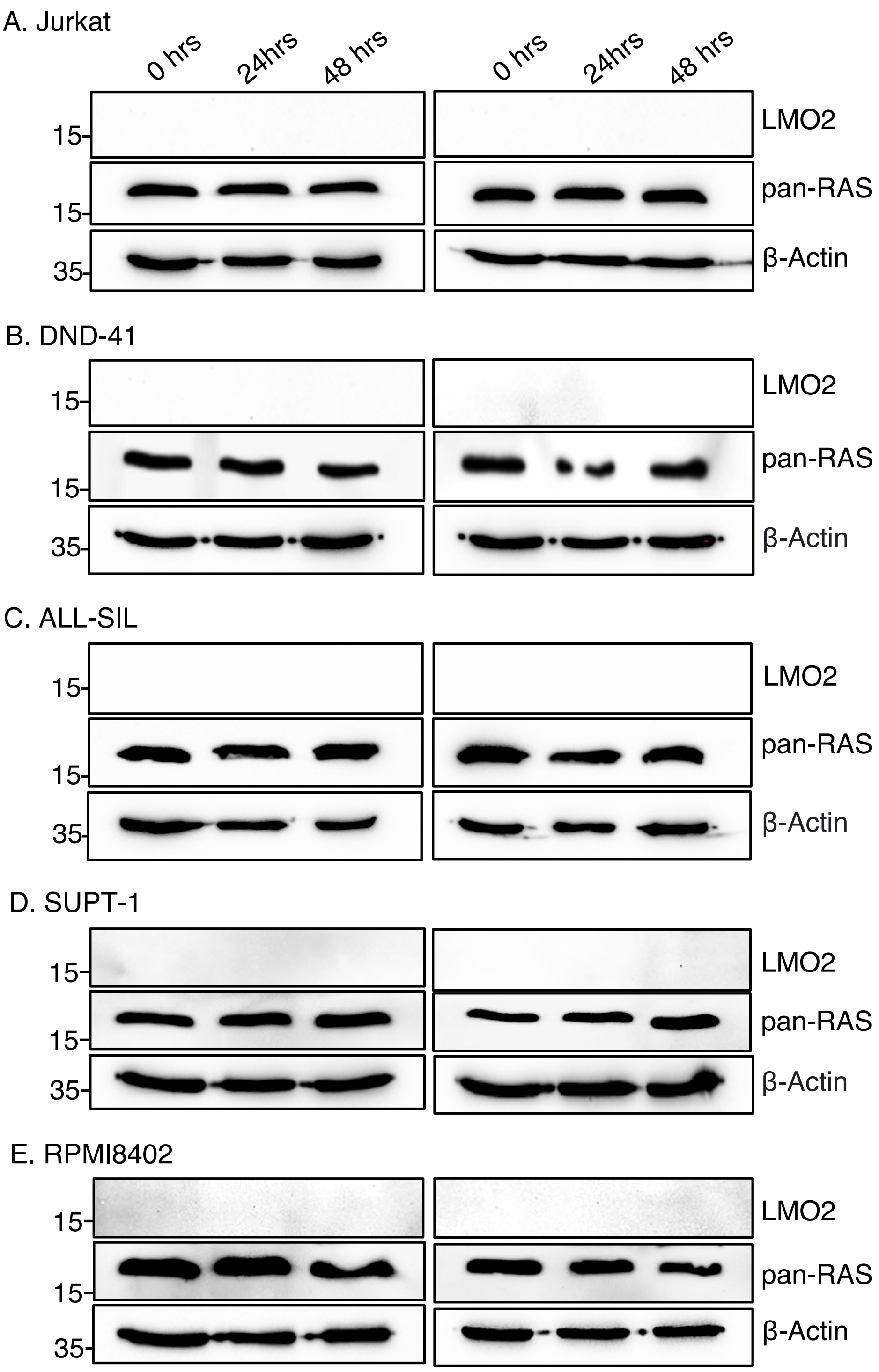


**Figure S7: Verification of LMO2 non-expressing T cell lines**

Five human T cell lines were verified by Western analysis of total protein extracts and probed with anti-LMO2, anti-RAS and β-actin as an internal loading control. No LMO2 was detectable. In addition, off-target effects, as determined by RAS protein levels, of the Abd-CRBN and Abd-VHL were tested by treating the cells with 20 µM Abd degraders for 24 hours and 48 hours followed by Western blotting.

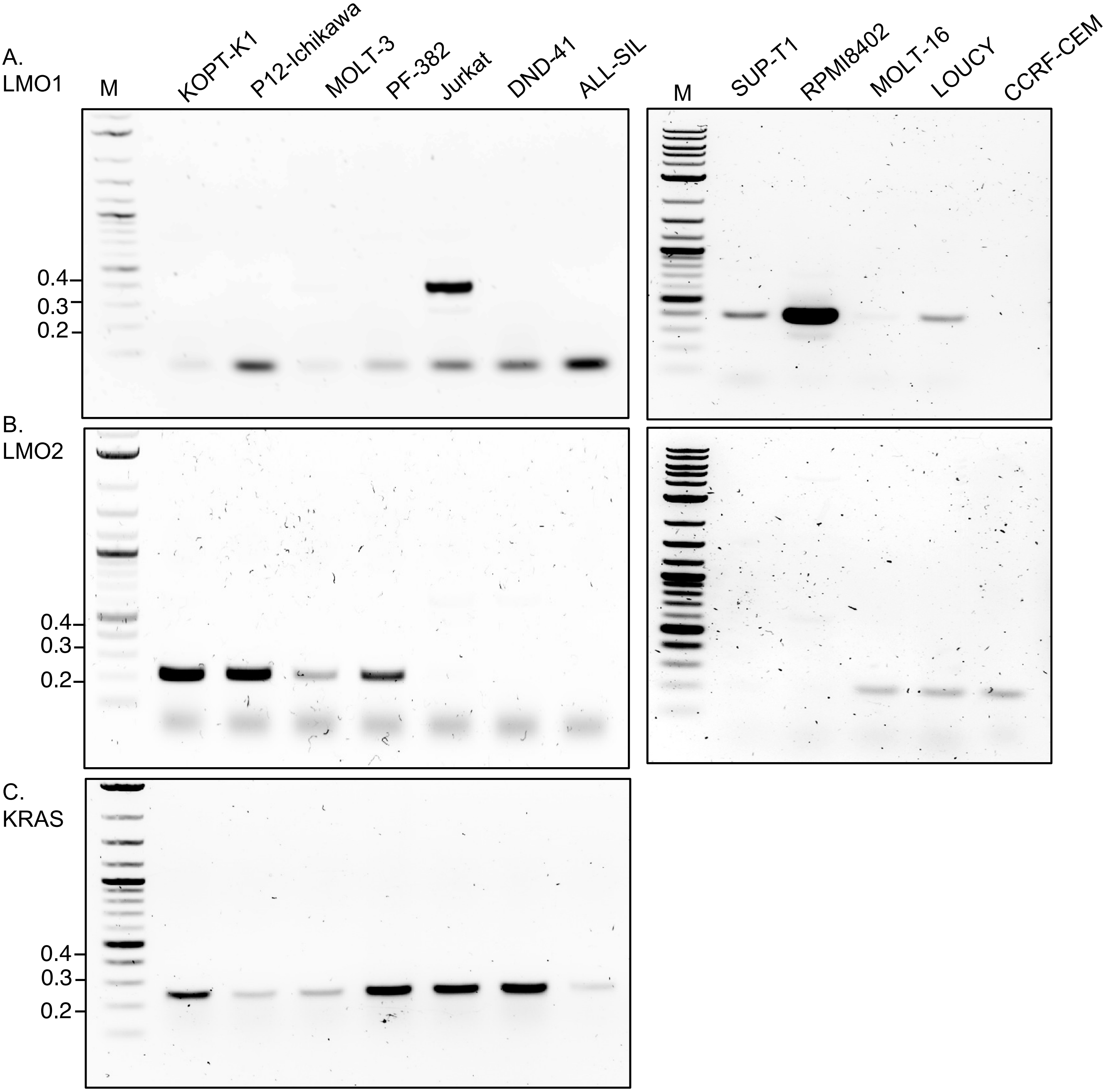


**Figure S8: RT-PCR analysis of the human T cell ALL panel**

Agarose gel electrophoresis of LMO1 (A), LMO2 (B), and KRAS (C) PCR products the different human T cell lines (KOPT-K1, P12-Ichikawa, MOLT-3, PF-382, Jurkat, DND-41, and ALL-SIL). Lane M represents a 1 kb ladder.

RT-PCR primer sequences.

LMO1: PCR product 410 bp

| | |
|---|---|
| Forward | GATCCAGCCCAAAGGGAAGCAG |
| Reverse | GATAAAGGTGCCATTGAGCTG |

LMO2: PCR product 220 bp

| | |
|---|---|
| Forward | GATTCCTCGGCCATCGAAAGG |
| Reverse | GATGTTTGTAGTAGAGGCGCCG |

KRAS: PCR product 249 bp

| | |
|---|---|
| Forward | GATATGACTGAATATAAACTTGTGGTAG |
| Reverse | GATGGCAAATACACAAAGAAAGC |

A.

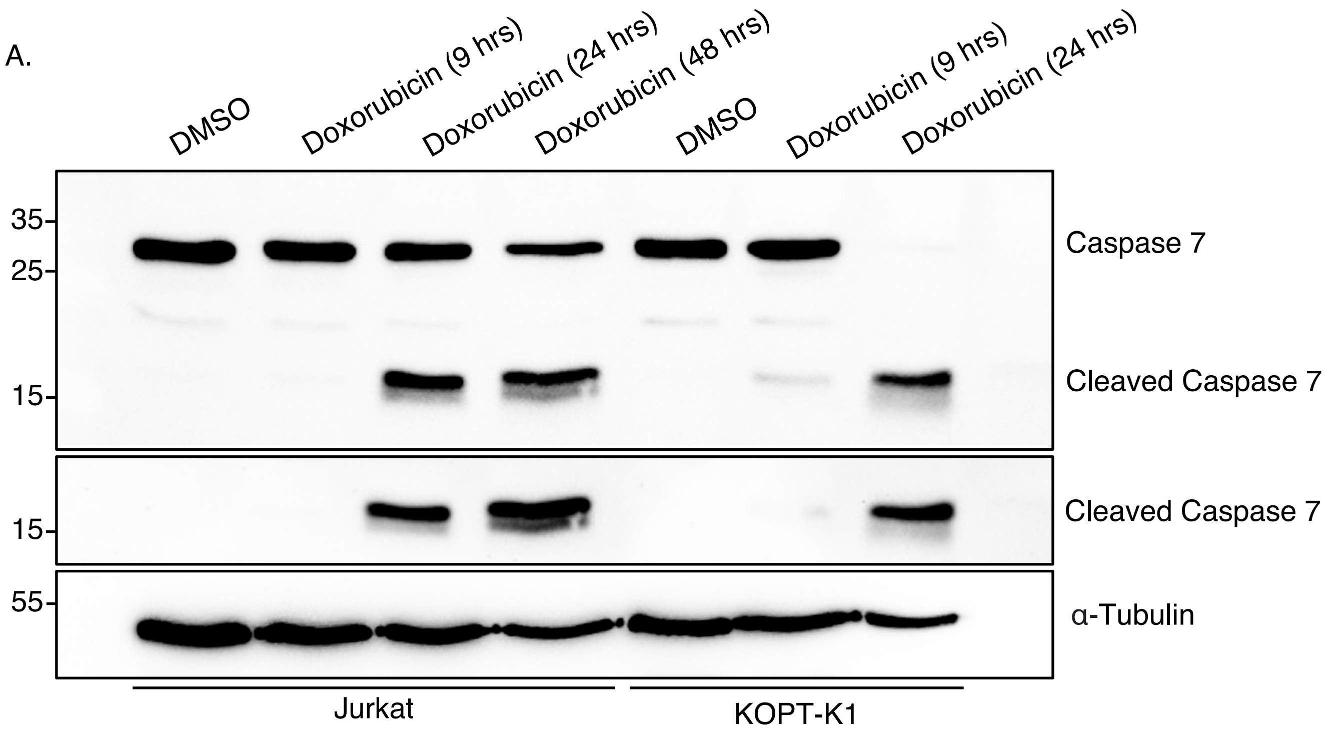


B.

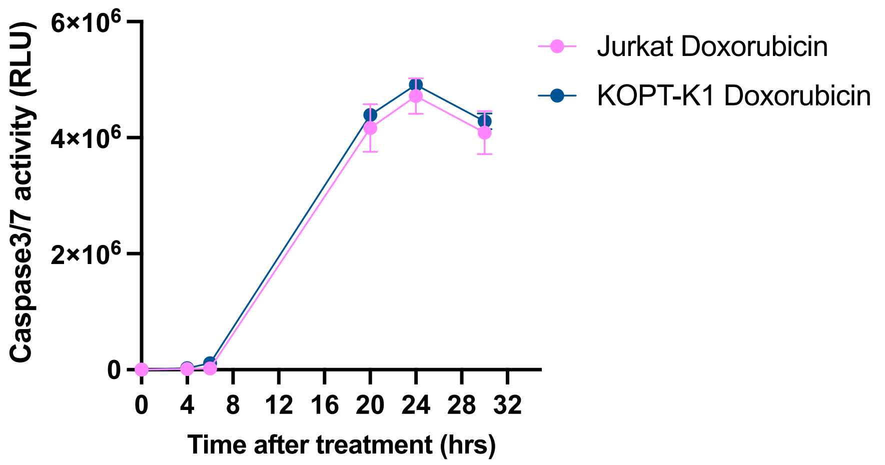


**Figure S9: Caspase7 and cleaved Caspase7 in Jurkat and KOPT-K1 cells after the treatment of doxorubicin.**

Western blot data showed Caspase7 and cleaved-Caspase7 after the treatment with 1 µM doxorubicin from 0 to 48 hours. At 48 hours treatment, doxorubicin at 1 µM induced 100% cell death in KOPT-K1 cells (panel A). Cell viability after the treatment at 4, 6, 12, 20, 24 and 30 hours in Jurkat and KOPT-K1 cells was determined using CellTiter-Glo assay (panel B). Data represent mean + SEM (n=3).

| Proteins | $log_2FC$ | Functions |
|---|---|---|
| AURKA | -0.376 | Kinase that contributes to the regulation of cell cycle progression |
| KIF2C | -0.384 | Important for anaphase chromosome segregation |
| HAUS1 | -0.394 | Maintenance of completion of cytokinesis |
| CDC25B | -0.428 | Activates the cyclin dependent kinase CDC2 |
| CCNA2 | -0.429 | Cyclin which controls both the G1/S and the G2/M transition |
| SGO2 | -0.466 | Part of mitotic cohesion complex |
| UBE2S | -0.482 | E2 enzyme ubiquitinates APC/C ubiquitin ligase |
| BUB1 | -0.488 | Member of the mitotic checkpoint complex and activating the spindle checkpoint. |
| USP37 | -0.575 | De-ubiquitinate enzyme promotes cell cycle |
| MASTL | -0.602 | MASTL kinase is a master regulator of mitosis |
| CDCA2 | -0.642 | Regulates chromosome structure during mitosis |
| UBE2C | -0.652 | E2 enzyme ubiquitinates APC/C |
| MIS18A | -0.704 | Required normal chromosome segregation during mitosis |
| FAM83D | -0.803 | Regulates cell proliferation, growth |
| CKS2 | -1.042 | Binds to CDKs for their function |
| TXNIP | -1.667 | Master regulator of cellular oxidation, regulating the expression and function of Thioredoxin |

**Figure S10: Proteins reduced by treatment of KOPT-K1 cells with Abd-VHL**
Proteomic data showed the significant downregulated proteins in KOPT-K1 cells when treated with Abd-VHL compared to control treatment (DMSO) and their functions related to cell division. Proteins shown based on the fold change ($log_2FC$) and p-value.

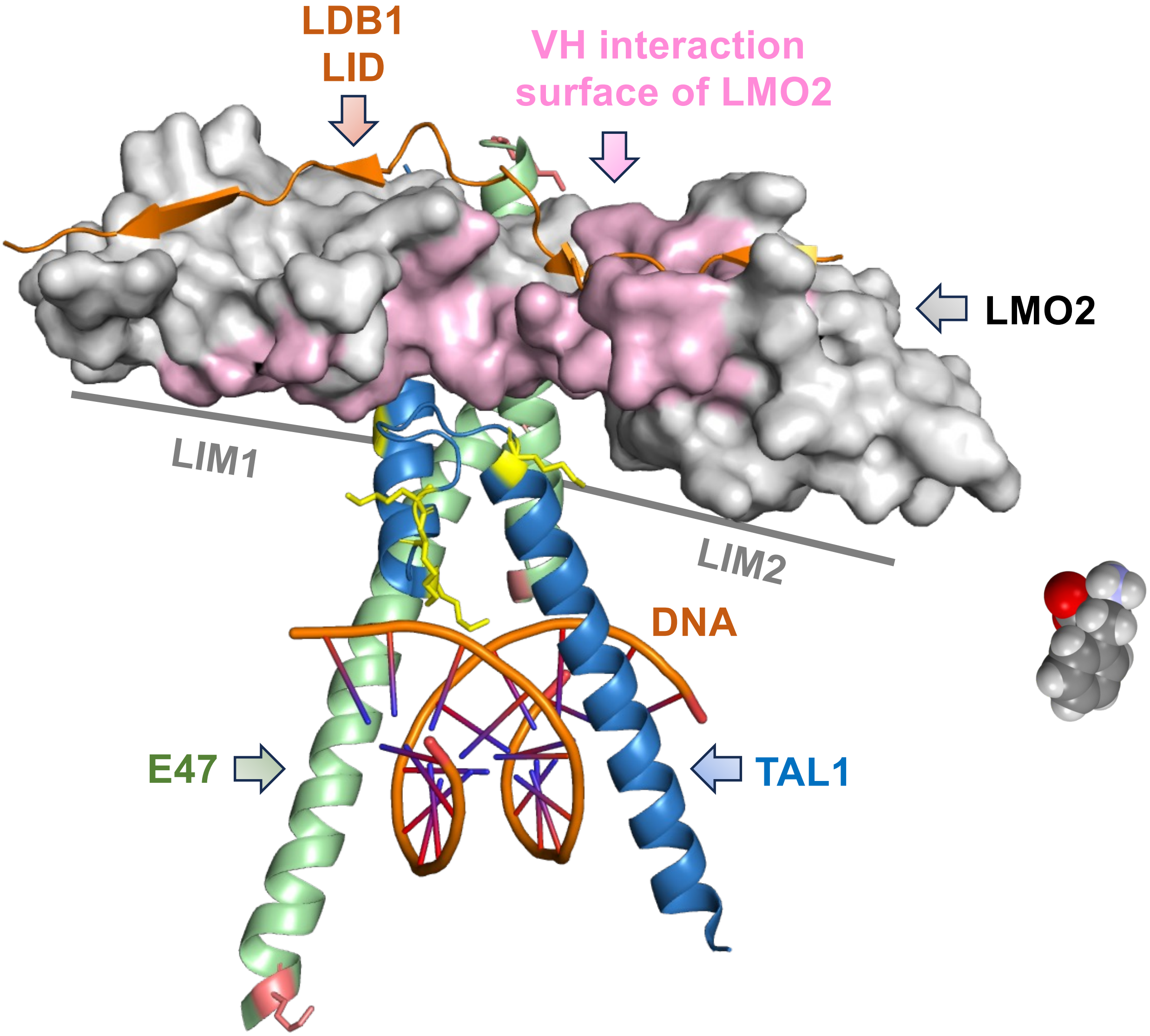


**Figure S11: The LMO2 transcription complex indicating lysine residues in the bHLH heterodimer for potential lysine ubiquitination**

The crystal structure of the LMO2-TAL1-E47 complex, adapted from (El Omari et al., 2013), is shown with LMO2 contacting the LID domain of LDB1, the TAL1 and E47 bHLH domains and DNA. LMO2 is shown in grey; the VH576 interaction domain in pink (Sewell et al., 2014); LID in orange; TAL1 in blue and E47 in green. Lysine residues in TAL1 and E47 are in yellow and salmon, respectively.

A.

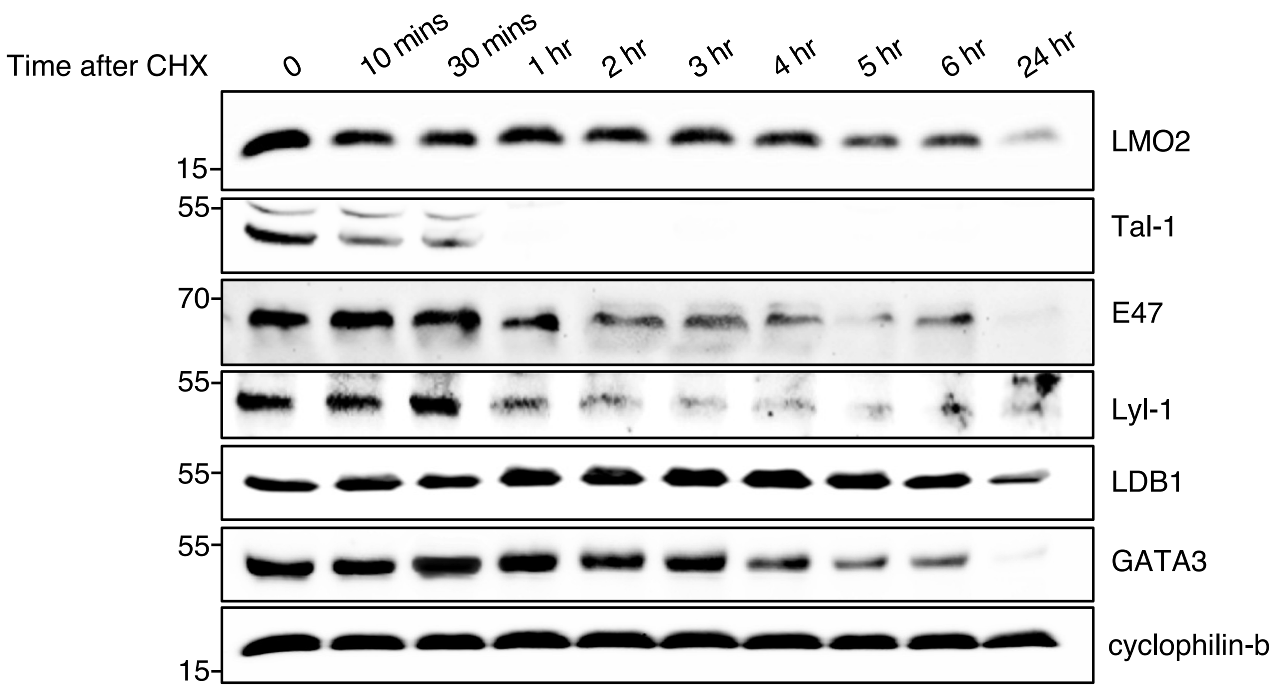


B.

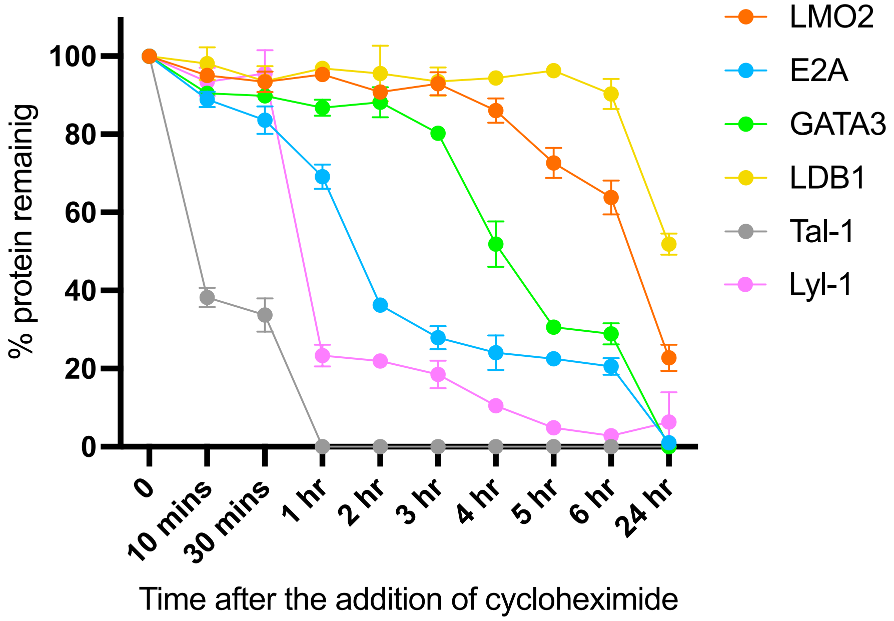


**Figure S12: Half-lives of LMO2 and LMO2 protein complex in KOPT-K1 cells.**
Western blot data showed LMO2 and LMO2 protein complex (TAL1, E47, LYL1, LDB1, and GATA3) level after the treatment with 20 µg/ml cycloheximide from 0 to 24 hours (panel A). The level of protein remaining presented as a relative to protein level at 0 hour (panel B). Data represent mean + SEM (n=3).

A. **gPCR primers**

KOPT FOR TWO(EcoRI)
GATGAATTCGAAGCTACTGCAGCCATC

KOPT FOR THREE (EcoRI)
GATGAATTCATGCTATGAGGTAGGTATG

J DELTA REV ONE (BamHI)
GATGGATCCGGTTCCACAGTCACTCGGGTTCC

B. **KOPT-K1 gPCR**

| | | |
|---|---|---|
| KOPT FOR TWO | + J DELTA REV ONE | 656bp |
| KOPT FOR THREE | + J DELTA REV ONE | 469bp |

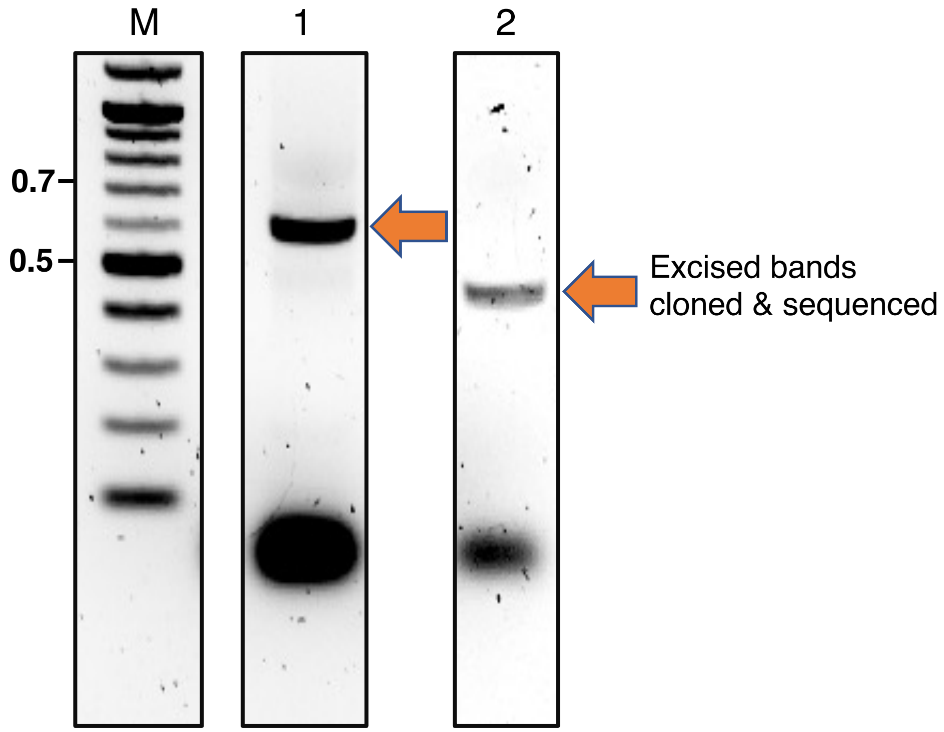


C.

```
GGAGGCTGGGGTAGGAGAATGAATTGAACCCAGGAGGCAGAGGTTTCAGTGAGCCAAGATCGGCCACTGCAT
TCCATCCTGGGTAACAGAGACTCCATCTCAATCAATCAATCAATCAATCAATAAAACAGTTTTATTGAGATG
TAATTGACATGCAATACGGCTACGGGGGGACGGGGATAACCAAAACTCATCTTTGGAAAAGGAACCCGAGTG
ACTGTGGAACCGGATCCACTAGTTCTAGAGCGGCCGCCACCGCGGTGGAGCTCCAGCTTTTGTTCCCTTTAG
TGAGGGTTAATTGCGCGCTTGGCGTAATCATGGTCATAGCTGTTTCCTGTGT
LMO2 Chr 11     B/P  Jdelta Chr 14
```

**Figure S13: Confirmation of chromosomal translocation in KOPT-K1 tissue culture cells**

The human T-ALL cells carries an LMO2 activating chromosomal translocation t(11;14)(p13;q11). The gPCR forward primers were designed to bind different 3'-end of the breakpoint region and reverse primer was designed to bind at 5'-end of the breakpoint region (panel A). Agarose gel electrophoresis was used to evaluate the PCR products (panel B), the different size of PCR products is showed in lane 1 and 2.
Lane M represents a 1 Kb marker ladder.
Panel C: The sequence of the genomic PCR primers were derived from the published breakpoint (Dong et al, 1995) and the relevant germline sequences of chromosomes 11 and 14.

**Supplementary Information**
**Synthesis of Abd-CRBN**

**ethyl 2-(4-(3-methoxybenzyl)piperazin-1-yl)oxazole-4-carboxylate**

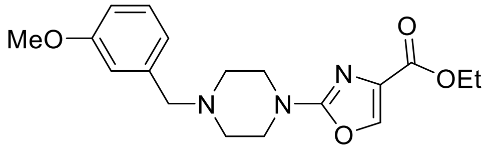


1-(3-Methoxybenzyl)piperazine (150 mg, 0.728 mmol, 1.2 eq.) was dissolved in 1,4-dioxane/*N*,*N*-diisopropylethylamine (4:1, 5 mL) before addition of ethyl 2-chlorooxazole-4-carboxylate (116 mg, 0.661 mmol, 1.0 eq.). The solution was stirred at 60 °C for 48 h, cooled down to room temperature, diluted with EtOAc (20 mL) and washed with $H_2O$/brine (1:1, 20 mL). The organic phase was dried ($Na_2SO_4$), filtered and concentrated *in vacuo*. The crude material was then purified on silica gel on silica gel (4% MeOH in $CH_2Cl_2$) afford the title compound as a a yellow oil (163 mg, 71%), after purification on silica gel (4% MeOH in $CH_2Cl_2$).

**ethyl 2-(4-(3-methoxybenzyl)piperazin-1-yl)oxazole-4-carboxylate**

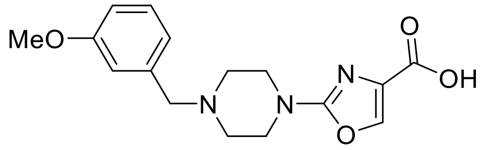


Ethyl 2-(4-(3-methoxybenzyl)piperazin-1-yl)oxazole-4-carboxylate (200 mg, 0.580 mmol, 1.0 eq.) was dissolved in THF/MeOH (4:1) before addition of NaOH (1M aq.) until pH> 8. The resulting reaction was stirred for 16 h at room temperature and then acidified with HCl (1M aq.) until pH<5. The solution was concentrated *in vacuo* and the obtained carboxylic acid (184 mg, quant.) used directly in the next step.

**ethyl 4-(2-(4-(3-methoxybenzyl)piperazin-1-yl)oxazole-4-carboxamido)benzoate**

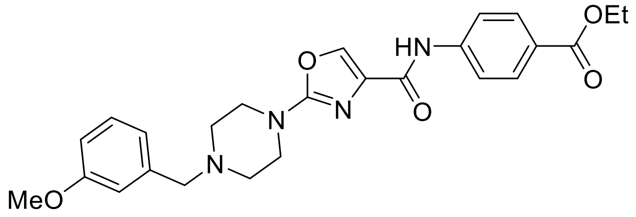


2-(4-(3-Methoxybenzyl)piperazin-1-yl)thiazole-4-carboxylic acid (184 mg, 0.580 mmol, 1.0 eq.) was dissolved in DMF (4 mL) before sequential addition of *N*,*N*-diisopropylethylamine (303 µL, 1.74 mmol, 3.0 eq.), benzocaine (115 mg, 0.696 mmol, 1.2 eq.) and HATU (309 mg, 0.812 mmol, 1.4 eq.). The resulting solution was stirred for 18 h, diluted with EtOAc (10 mL) and washed with brine/water (1:1, 3 x 50 mL). The organic phase was dried ($Na_2SO_4$), filtered and concentrated *in vacuo*. The title product was obtained as a colourless oil that solidified on standing (247 mg, 92%), after purification on silica gel (4% MeOH in $CH_2Cl_2$).

**4-(2-(4-(3-methoxybenzyl)piperazin-1-yl)oxazole-4-carboxamido)benzoic acid**

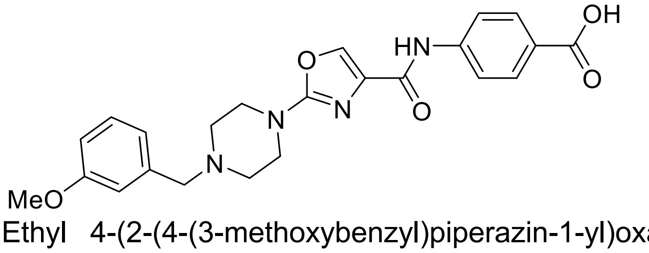


Ethyl 4-(2-(4-(3-methoxybenzyl)piperazin-1-yl)oxazole-4-carboxamido)benzoate (120 mg, 0.252 mmol, 1.0 eq.) was dissolved in THF/MeOH (4:1) before addition of NaOH (1M aq.) until pH> 8. The resulting reaction was stirred for 16 h at room temperature and then acidified with

HCl (1M aq.) until pH<5. The solution was concentrated *in vacuo* and the obtained carboxylic acid (114 mg, quant.) used directly in the next step.

**_N_-(4-((6-((2-(2,6-dioxopiperidin-3-yl)-1,3-dioxoisoindolin-4-yl)oxy)hexyl)carbamoyl)phenyl)-2-(4-(3-methoxybenzyl)piperazin-1-yl)oxazole-4-carboxamide**

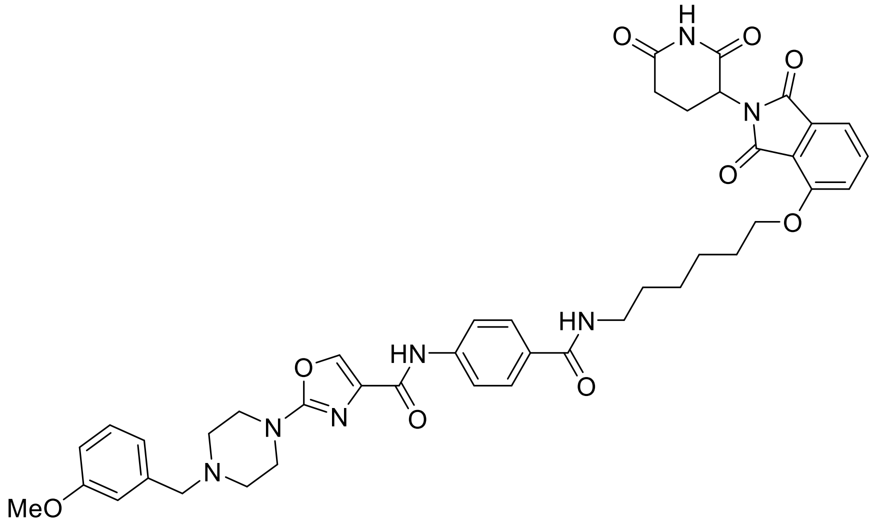


4-(2-(4-(3-Methoxybenzyl)piperazin-1-yl)oxazole-4-carboxamido)benzoic (65 mg, 0.150 mmol, 1.2 eq.) was dissolved in DMF (1 mL) before sequential addition of *N*,*N*-diisopropylethylamine (66 µL, 0.381 mmol, 3.0 eq.), 4-((6-aminohexyl)oxy)-2-(2,6-dioxopiperidin-3-yl)isoindoline-1,3-dione (47 mg, 0.127 mmol, 1.0 eq.) and HATU (80 mg, 0.210 mmol, 1.4 eq.). The resulting solution was stirred for 18 h, diluted with EtOAc (10 mL) and washed with brine/water (1:1, 3 x 50 mL). The organic phase was dried ($Na_2SO_4$), filtered and concentrated *in vacuo*. The title product was obtained as a colourless oil (89 mg, 89%), after purification on silica gel (10% MeOH in $CH_2Cl_2$).

#### Synthesis of Abd-VHL

*tert*-Butyl 2-(2-(2-aminoethoxy)ethoxy)acetate was prepared followed literature precedent.

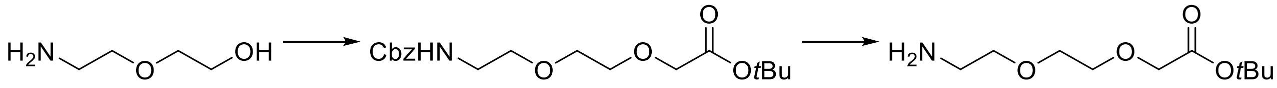

##### ***tert*-butyl 2-(2-(2-(4-(2-(4-(3-methoxybenzyl)piperazin-1-yl)thiazole-4-carboxamido) benzamido)ethoxy)ethoxy)acetate**

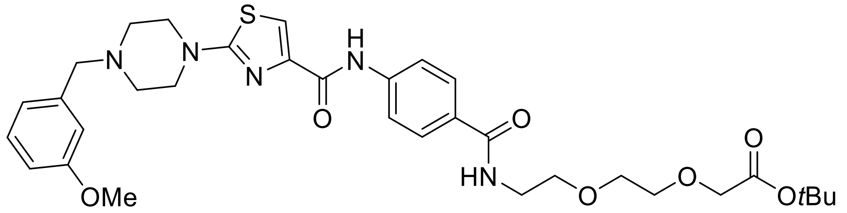

4-(2-(4-(3-Methoxybenzyl)piperazin-1-yl)thiazole-4-carboxamido)benzoic (36 mg, 0.080 mmol, 1.0 eq.) was dissolved in DMF (1 mL) before sequential addition of *N*,*N*-diisopropylethylamine (42 µL, 0.240 mmol, 3.0 eq.), *tert*-butyl 2-(2-(2-aminoethoxy)ethoxy)acetate (20 mg, 0.086 mmol, 1.2 eq.) and HATU (43 mg, 0.112 mmol, 1.4 eq.). The resulting solution was stirred for 18 h, diluted with EtOAc (10 mL) and washed with brine/water (1:1, 3 x 50 mL). The organic phase was dried ($Na_2SO_4$), filtered and concentrated *in vacuo*. The title product was obtained as a colourless oil (30 mg, 57%), after purification on silica gel (10% MeOH in $CH_2Cl_2$).

##### **2-(2-(2-(4-(2-(4-(3-methoxybenzyl)piperazin-1-yl)thiazole-4 carboxamido)benzamido) ethoxy)ethoxy) acetic acid**

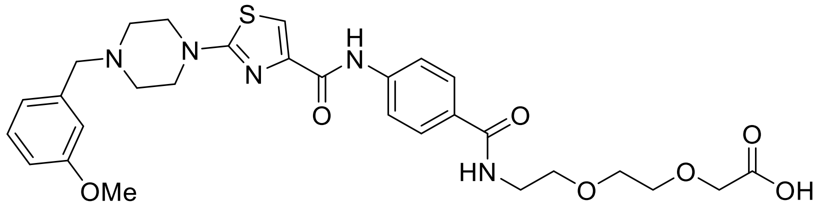

*tert*-Butyl 2-(2-(2-(4-(2-(4-(3-methoxybenzyl)piperazin-1-yl)thiazole-4-carboxamido) benzamido)ethoxy)ethoxy)acetate (30 mg, 0.046 mmol, 1.0 eq.) was dissolved in $CH_2Cl_2$ (1 mL) before addition of TFA (10 µL) in one portion. The reaction was stirred for 16 h at room temperature, concentrated in vacuo and the title product was obtained as a yellow oil (27 mg, quant.) and used directly in the next step.

##### **VHL ligand:**

The VHL ligand ((2*S*,4*R*)-1-((*R*)-2-amino-3,3-dimethylbutanoyl)-4-hydroxy-*N*-(4-(4-methylthiazol-5-yl)benzyl)pyrrolidine-2-carboxamide) was prepared following literature precedent with a small modification; the first 2 steps were replaced by a Suzuki reaction. The final product was columned before use (not carried out in the literature).

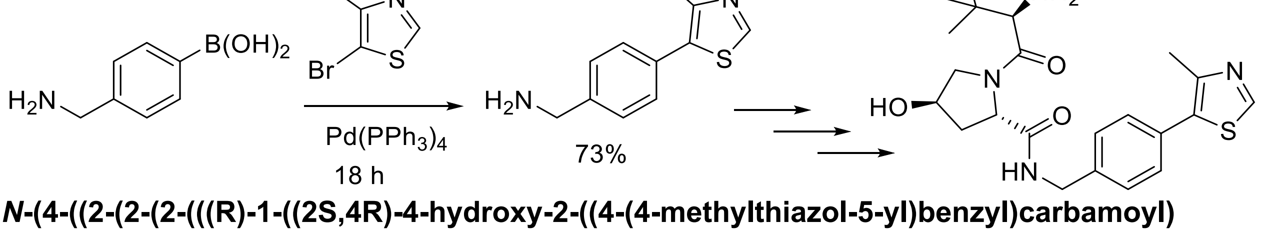


##### ***N*-(4-((2-(2-(2-(((R)-1-((2S,4R)-4-hydroxy-2-((4-(4-methylthiazol-5-yl)benzyl)carbamoyl) pyrrolidine-1-yl)-3,3-dimethyl-1-oxobutan-2-yl)amino)-2-oxoethoxy)ethoxy)ethyl) carbamoyl)phenyl)-2-(4-(3-methoxybenzyl)piperazin-1-yl)thiazole-4-carboxamide**

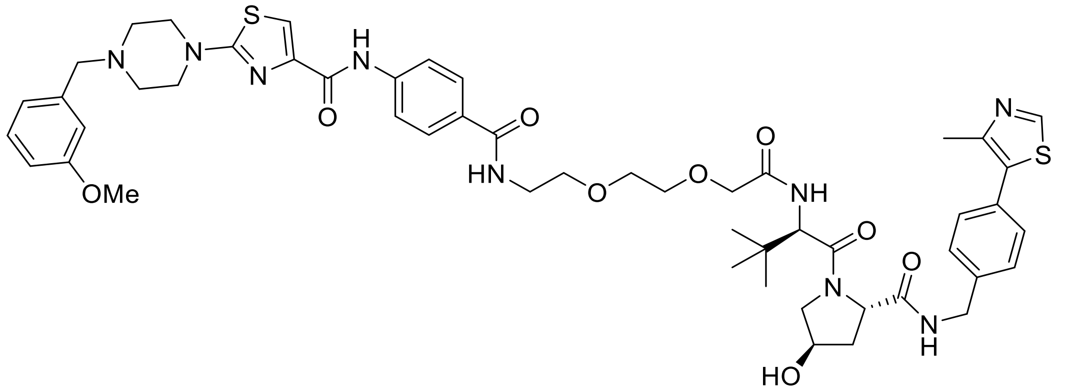

2-(2-(2-(4-(2-(4-(3-Methoxybenzyl)piperazin-1-yl)thiazole-4 carboxamido)benzamido) ethoxy)ethoxy) acetic acid (20 mg, 0.033 mmol, 1.0 eq.) was dissolved in DMF (1 mL) before sequential addition of *N*,*N*-diisopropylethylamine (17 µL, 0.100 mmol, 3.0 eq.), 2*S*,4*R*)-1-((*R*)-2-amino-3,3-dimethylbutanoyl)-4-hydroxy-*N*-(4-(4-methylthiazol-5-yl)benzyl)pyrrolidine-2-carboxamide (17 mg, 0.040 mmol, 1.2 eq.) and HATU (18 mg, 0.046 mmol, 1.4 eq.). The resulting solution was stirred for 18 h, diluted with EtOAc (10 mL) and washed with brine/water (1:1, 3 x 50 mL). The organic phase was dried ($Na_2SO_4$), filtered and concentrated *in vacuo*. The title product was obtained as a colourless oil (9 mg, 27%), after purification on silica gel (18% MeOH in $CH_2Cl_2$), followed by two successive preparative TLC (15% MeOH in $CH_2Cl_2$ with double elution).
